# Oviposition of the mosquito *Aedes aegypti* in forest and domestic habitats in Africa

**DOI:** 10.1101/2020.07.08.192187

**Authors:** Siyang Xia, Hany K. M. Dweck, Joel Lutomiah, Rosemary Sang, Carolyn S. McBride, Noah H. Rose, Diego Ayala, Jeffrey R. Powell

## Abstract

The theory of ecological divergence provides a useful framework to understand the adaptation of many species to anthropogenic (‘domestic’) habitats. The mosquito *Aedes aegypti*, a global vector of several arboviral diseases, presents an excellent study system. *Ae. aegypti* originated in African forests, but the populations that invaded other continents have specialized in domestic habitats. In its African native range, the species can be found in both forest and domestic habitats like villages. A crucial behavioral change between mosquitoes living in different habitats is their oviposition choices. Forest *Ae. aegypti* lay eggs in natural water containers like tree holes, while their domestic counterparts heavily rely on artificial containers such as plastic buckets. These habitat-specific containers likely have different environmental conditions, which could drive the incipient divergent evolution of oviposition in African *Ae. aegypti*. To examine this hypothesis, we conducted field research in two African locations, La Lopé, Gabon and Rabai, Kenya, where *Ae. aegypti* live in both forests and nearby villages. We first characterized a series of environmental conditions of natural oviposition sites, including physical characteristics, microbial density, bacterial composition, and volatile profiles. Our data showed that in both locations, environmental conditions of oviposition sites did differ between habitats. To examine potential behavioral divergence, we then conducted field and laboratory oviposition choice experiments to compare the oviposition preference of forest and village mosquitoes. The field experiment suggested that forest mosquitoes readily accepted artificial containers. In laboratory oviposition assays, forest and village mosquito colonies did not show a differential preference towards several conditions that featured forest versus village oviposition sites. Collectively, there is little evidence from our study that environmental differences lead to strong and easily measurable divergence in oviposition behavior between *Ae. aegypti* that occupy nearby forest and domestic habitats within Africa, despite clear divergence between African and non-African *Ae. aegypti*.

## Introduction

Ecological divergence is one of the central mechanisms contributing to biodiversity (Nosil, 2012). When descendants of the same ancestral population evolve in different environments, they may experience divergent selection pressures leading to morphological and/or behavioral divergence (Schluter, 2000). Accumulation of these phenotypic changes and their underlying genetic components, along with genetic drift, could further result in reproductive isolation and speciation (Nosil, 2012; Rundle & Nosil, 2005; Shafer & Wolf, 2013). A core step in this process is the ecologically-based divergent selection (Rundle & Nosil, 2005), which can be recognized by two essential features: consistently distinct environmental conditions and organisms’ corresponding phenotypic adaptations. The attribution of phenotypic change to ecological selection has been demonstrated in several natural populations, such as Darwin’s finches (Grant & Grant, 2002, 2011), stickleback fish (Hatfield & Schluter, 1999), beach mice (Mullen, Vignieri, Gore, & Hoekstra, 2009), and *Timema cristinae* walking-sticks (Nosil, 2007; Nosil & Crespi, 2004).

In addition to explaining biodiversity, the model of ecological divergence also provides a useful framework for understanding the evolution of a particular group of organisms – disease vectors living with humans, such as mosquitoes. Many of these vector species experienced a transition from their natural habitats into anthropogenic domestic habitats (e.g., villages and urban areas) following the development of human civilization (Hulme-Beaman, Dobney, Cucchi, & Searle, 2016; Otto, 2018). The striking contrast between these two types of habitats suggests a potentially strong divergent selection (Johnson & Munshi-South, 2017). Alternatively, some vectors species may be predisposed to using domestic habitats, in which case one would expect little phenotypic changes. Few studies have examined these hypotheses or demonstrated how these disease vector species react and adapt to the ‘novel’ environmental conditions of the domestic habitats. Addressing this question will contribute to our understanding of the unique evolutionary history of these epidemiologically important animals, and provide valuable information on why they are so good at living around humans and transmitting diseases.

The mosquito *Aedes aegypti* provides an excellent model for studying ecological divergence in disease vectors. The species is the main vector of yellow fever, dengue, chikungunya (World Health Organization, 2014), and Zika virus (Li, Wong, Ng, & Tan, 2012; Marcondes & Ximenes, 2016). Genetic data suggested that *Ae. aegypti* is native to Africa (Brown et al., 2011; Gloria-Soria et al., 2016; Powell, Gloria-Soria, & Kotsakiozi, 2018). With the establishment of human settlements, they invaded human-generated domestic habitats inside Africa, probably five to ten thousand years ago (Crawford et al., 2017; Kotsakiozi, Evans, et al., 2018), and later spread to the rest of the world since the 15^th^ century (Brown et al., 2014; Powell et al., 2018; Powell & Tabachnick, 2013). The mosquitoes in and out of Africa showed a relatively clear genetic distinction (but see exceptions in Kotsakiozi et al. 2018 and Rose et al. 2020), which roughly matches the two classical subspecies: *Ae. aegypti formosus* (Aaf) and *Ae. aegypti aegypti* (Aaa), respectively. Complexities exist in this subspecies definition (Powell & Tabachnick, 2013), and in this paper, we refer to them simply based on their geographic range (in or out of Africa). Non-African Aaa breed in human environments, e.g., live specifically in urban areas with only a few exceptions in the Caribbean and Argentina (Chadee, Ward, & Novak, 1998; Mangudo, Aparicio, & Gleiser, 2015). They also display a strong preference for human hosts (McBride et al., 2014; Rose et al., 2020) and use artificial containers as breedings sites (Day, 2016).

However, relatively little is known about the initial process of colonizing domestic environments inside Africa, except for a few recent studies. For example, Rose et al. (2020) found that the mosquito’s preference towards humans is closely associated with seasonality and human density. *Ae. aegypti* throughout Africa (Aaf) can be found in both forests, the presumed ancestral habitats, and domestic settings like villages. Previous studies showed that domestic Aaf in several locations inside Africa is genetically similar to their local forest counterparts, suggesting a relatively recent invasion into domestic habitats (Kotsakiozi, Evans, et al., 2018; Paupy et al., 2014; Sylla, Bosio, Urdaneta-Marquez, Ndiaye, & Black IV, 2009). Comparing Aaf between different habitats could allow us to understand the potential incipient divergence. For example, what were the original selective pressures that may have ultimately led to the clear divergence between Aaf and Aaa?

From a behavioral perspective, one of the critical steps during the process of colonizing domestic habitats and invading other tropical regions around the world is adapting to lay eggs (i.e., oviposit) in domestic breeding sites. After taking a full blood meal, which is necessary for reproduction, *Ae. aegypti* females lay eggs on substrates at the edge of small containers of water, i.e., oviposition sites (Christophers, 1960). Aaf in African forest and domestic habitats utilize different oviposition sites: the former lay eggs mainly in natural containers like water-filled tree holes and rock pools (Lounibos, 1981), while the latter uses mostly artificial containers, such as plastic buckets, tires, and discarded tin cans (McBride et al., 2014; Petersen, 1977; Trpis & Hausermann, 1978). This difference in oviposition site use is at least partly a function of container availability in the two habitats. However, natural and artificial containers likely have different characteristics (Yee, Allgood, Kneitel, & Kuehn, 2012), such as bacterial profiles (Dickson et al., 2017), that could also drive genetically based divergence in container preference. Such divergence likely exists between Aaa and Aaf, as shown in studies comparing Aaf and a once existed Aaa introduced to coastal Kenya from non-African populations (Leahy, VandeHey, & Booth, 1978; Petersen, 1977). Whether a similar divergence also exists within Aaf remains mostly unclear.

Conversely, ovipositional modifications could have a significant effect on the evolution of the mosquitoes. If forest and domestic Aaf actively prefer natural and artificial containers, respectively, it could facilitate the isolation between them: selective oviposition could keep forest populations in the forest and domestic populations close to humans, which reduces gene flow between them and promotes other adaptations (Servedio, Van Doorn, Kopp, Frame, & Nosil, 2011). Therefore, the evolution of oviposition behaviors could be a key factor in understanding how *Ae. aegypti* became domesticated (Powell et al., 2018; Rose et al., 2020). This process is of particular interest in the initial colonization of domestic habitat within Africa. How different are the oviposition sites in the forest versus domestic habitats? Was Aaf an ovipositional generalist, pre-adapted to jump into human environments? Or are there genetically-based differences between populations breeding in wild and human environments?

*Ae. aegypti* choose oviposition sites based on the interactions between their innate preference and external oviposition cues. Various abiotic and biotic factors have been shown to influence oviposition choices of *Ae. aegypti* (Day, 2016), including container size (Bond & Fay, 1969; Burkot et al., 2007; Harrington, Ponlawat, Edman, Scott, & Vermeylen, 2008), shading (Barrera, Amador, & Clark, 2006; Prado, Maciel, Leite, & Souza, 2017), water salinity (Matthews, Younger, & Vosshall, 2019), color and texture of the sites (Bentley & Day, 1989; Fay & Perry, 1965), presence of conspecific eggs, larvae, and pupae (Zahiri, Rau, & Lewis, 1997, 1997), predators (Albeny-Simoes et al., 2014; Pamplona Lde, Alencar, Lima, & Heukelbach, 2009), bacterial density and community composition (Arbaoui & Chua, 2014; Hazard, Mayer, & Savage, 1967; Ponnusamy, Schal, Wesson, Arellano, & Apperson, 2015), and chemical components (Afify & Galizia, 2015; Melo et al., 2019). However, most of the existing studies were conducted in laboratory settings with artificial oviposition choices. Although these studies provided rich knowledge on the sensory mechanisms of oviposition (Matthews et al., 2019; Ponnusamy et al., 2015), the conditions examined in these studies may not necessarily reflect the characteristics of breeding sites in the field. These studies are also heavily biased to Aaa, while detailed examination of oviposition preference in Aaf is mostly missing, let alone comparisons between forest and domestic Aaf. As a result, it is still unclear how oviposition behaviors evolved during the domestication of *Ae. aegypti*.

As a first step to address this question, we examined oviposition of *Ae. aegypti* living in forest and domestic habitats in two locations in Africa, La Lopé in Gabon and Rabai in Kenya. Mosquitoes in both locations are Aaf, but can be found in forest and villages in close proximity. They also showed little genetic differentiation between habitats (Xia et al., submitted), which suggested gene flow between forest and domestic populations. We first characterized the environmental conditions of natural oviposition sites, including physical charasteristics, competition and predation, bacterial profiles, and chemical volatiles, in natural sites (tree holes) and artificial containers. To examine whether environmental differences may translate into behavioral differences, we then investigated the oviposition preference of forest and domestic Aaf through field oviposition experiments and laboratory oviposition assays. The results could also provide useful information on identifying the critical environmental variables that potentially drove the divergent evolution of oviposition, if such behavioral divergence exists.

We hypothesized that natural and artificial containers represent very different environmental characteristics, and that both forest and domestic Aaf will prefer conditions that are more alike the oviposition sites from their own habitats, as would be predicted under a model of ecological divergence and local adaptation. By examining the two main elements of ecological divergence, environmental variation and behavioral differences, this study provides valuable information on how oviposition behaviors in *Ae. aegypti* evolved during the domestication history of the mosquito.

## Materials and methods

### Field study

We conducted field studies in La Lopé, Gabon in Central Africa from November to December 2016, and in Rabai, Kenya in East Africa from April to May 2017. La Lopé has an extensive continuous tropical rainforest surrounding La Lopé village (Figure 1a). The forest in Rabai, on the other hand, is more fragmented, with several villages scattered around the forest patch (Figure 1b). In each location, we searched for water-holding containers as potential mosquito oviposition sites in both the forests and nearby villages. A potential oviposition site was defined as one that holds at least one mosquito larva (not necessarily *Ae. aegypti*) at the time of sampling, which suggested that the site had been present long enough for a mosquito to lay eggs. We categorized oviposition sites into three habitat groups: forest, peridomestic (outdoor containers in a village area), and domestic (indoor containers) (Table 1). We separated indoor and outdoor containers as classical studies from the 1970s reported that, at least in Rabai, Kenya, the mosquitoes living indoor and outdoor showed distinct behavioral and genetic difference (Leahy et al., 1978; McBride et al., 2014; Petersen, 1977; Tabachnick, Munstermann, & Powell, 1979; Trpis & Hausermann, 1975). Genetic analysis showed that these indoor mosquitoes in Rabai were likely descendent of non-African Aaa, which is a unique case in the evolutionary history of *Ae. aegypti* (Brown et al., 2011; Gloria-Soria et al., 2016). However, this previously described Aaa-like indoor form was no longer found at the time of sampling, which is also supported by genetic data (Xia et al., submitted).

**Figure 1.**
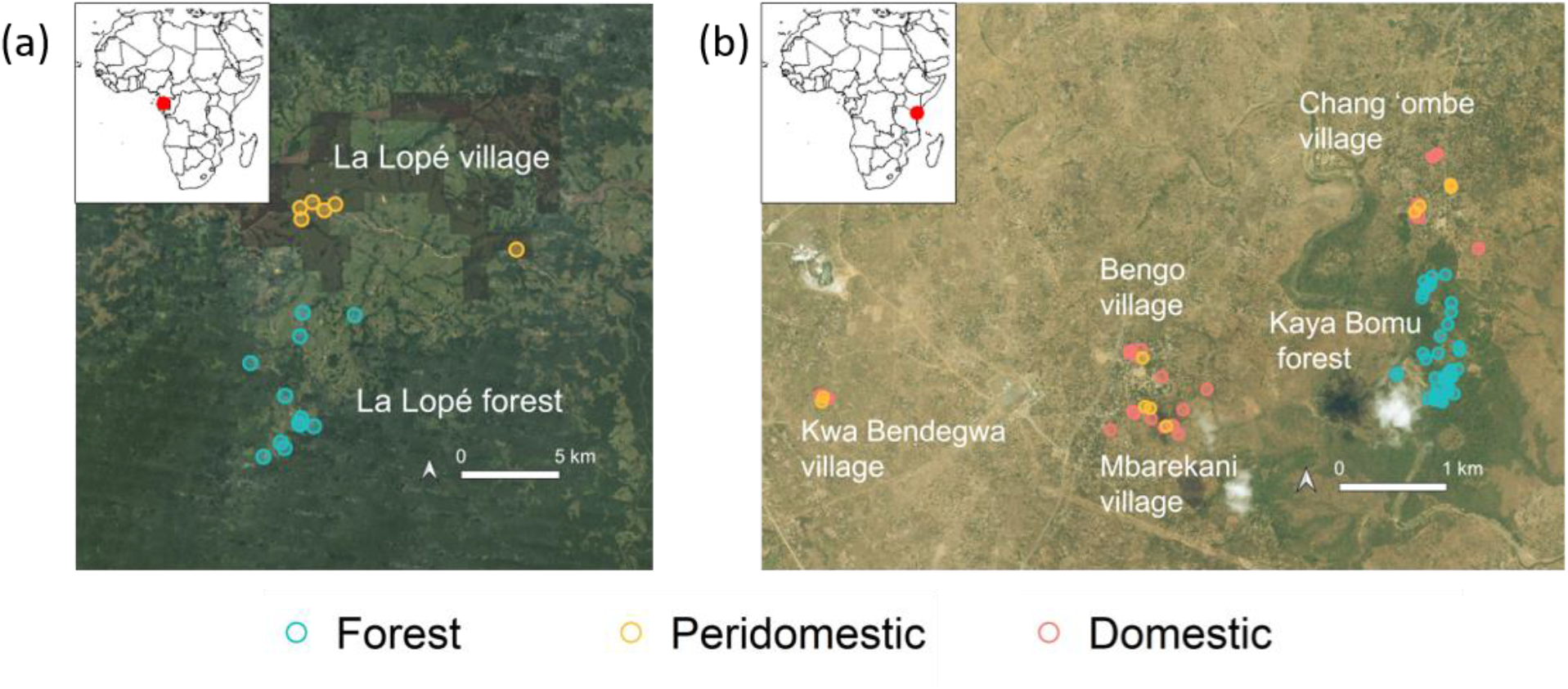
Sampling locations in (a) La Lopé, Gabon, and (b) Rabai, Kenya. The inset in each graph shows the location of the field site in continental Africa. In (a), each point represents a sampling site where one to multiple oviposition sites were found. In (b), each point represents a single oviposition site. The color of the point indicates the habitat category: red points are domestic (village indoor) sites, yellow points are peridomestic sites (village outdoor), and green points are forest sites. The satellite image were from (a) Google Satellite and (b) Bing Satellite in QGIS, respectively.

**Table 1.** Number of oviposition sites measured for different environmental variables

| Field site location | Habitat | *Aedes aegypti* | Physical characteristics | Larval density | Microbial density | Bacteria composition | Volatile profile |
| --- | --- | --- | --- | --- | --- | --- | --- |
| La Lopé,<br>Gabon | Forest | Present | 5 | 5 | 5 | 5 | na<sup>*</sup> |
|  |  | Absent | 48 | 55 | 10 | 33 | na<sup>*</sup> |
|  | Peridomestic<br>(Village) | Present | 13 | 13 | 10 | 10 | na<sup>*</sup> |
|  |  | Absent | 24 | 25 | 12 | 23 | na<sup>*</sup> |
|  | **Total** |  | 90 | 98 | 37 | 71 | na<sup>*</sup> |
| Rabai,<br>Kenya | Forest | Present | 15 | 15 | 15 | 15 | 7 |
|  |  | Absent | 22 | 22 | 11 | 22 | 12 |
|  | Peridomestic<br>(Village) | Present | 8 | 8 | 8 | 8 | 5 |
|  |  | Absent | 1 | 1 | 1 | 1 | 1 |
|  | Domestic<br>(Village) | Present | 22 | 22 | 22 | 22 | 17 |
|  | **Total** |  | 68 | 68 | 57 | 68 | 42 |

<sup>\*</sup> Headspace volatiles were not collected in La Lopé, Gabon.

In La Lopé, we visited 60 oviposition sites in seven forest locations, and 38 sites in six village locations. The sampling sites separate by 5-17 km. Forest oviposition sites were predominantly rock pools around streams and tree holes that accumulated water. In the village, mosquito larvae were found in a variety of artificial containers, including construction bricks, tires, metal cans, and plastic containers. Because residents in the village rarely store water indoor, all village oviposition sites were ‘peridomestic.’ In Rabai, Kenya, we sampled 31 oviposition sites consisting of mainly plastic buckets, earthenware pots, and metal barrels in four villages. They were mostly indoor (i.e., domestic) containers. The 37 oviposition sites in Rabai forest were all tree holes holding rainwater (Figure 1b). We recorded the GPS coordinates of each sampling location (which may consist of more than one oviposition sites) in La Lopé, and of each oviposition site in Rabai, Kenya (Figure 1).

Upon identifying a potential oviposition site in any habitat, we measured 11-16 physical variables. We also collected water samples for further analysis of bacterial and chemical volatile profiles. Method details are described in the following sections and the Appendix. In addition, we collected all mosquito larvae using pipets and reared them to adults in field stations, keeping larvae and pupae from different oviposition sites separate. Upon eclosion, the adults were identified to species or genus based on taxonomic keys using a dissection microscope in the field. We kept *Ae. aegypti* adults alive to establish lab colonies for future behavioral tests. We categorized each site as ‘*Ae. aegypti* present’ or ‘*Ae. aegypti* absent’ based on whether it held any *Ae. aegypti* larvae or pupae (Table 1). It is worth noting that the absence of *Ae. aegypti* may not necessarily suggest an avoidance. Some sites may be suitable for oviposition but not yet colonized by *Ae. aegypti* at the time of collection. The combinations of habitats and *Ae. aegypti* presence will be referred to as ‘oviposition site groups’ in the rest of the paper for the purpose of communication. We summarized the sample sizes for analyses of different environmental variables in Table 1. Almost all peridomestic and domestic habitats in Rabai were present with *Ae. aegypti*. This mainly results from the fact that there were rarely other species present in these environments, but we required at least one mosquito larvae to include the site in the dataset, so there are effectively no ‘*Ae aegypti* absent’ sites. Because the peridomestic *Ae. aegypti* absent group contained only one sample, it is excluded from group-level analyses, but retained in comparisons between habitats or between *Ae. aegypti* present vs. absent sites.

The fieldwork in La Lopé was approved by the CENAREST with the authorization AR0013/16/MESRS/CENAREST/CG/CST/CSAR, and by the La Lopé National Parks with the authorization AE16008/PR/ANPN/SE/CS/AEPN. The fieldwork in Rabai was approved by the Kenya Medical Research Institute Scientific and Ethical Review Unit with the authorization KEMRI/SERU/3433.

### Characterizing oviposition sites: physical variables

We measured 11 physical variables for each oviposition site in La Lopé, Gabon, and five additional variables in Rabai, Kenya (Table S1 in Appendix). The variables were selected partially based on previous laboratory studies of mosquito oviposition (Harrington et al., 2008; Madeira, Macharelli, & Carvalho, 2002; Petersen, 1977; Reiskind & Zarrabi, 2012; Wong, Stoddard, Astete, Morrison, & Scott, 2011), as well as the availability of equipment and resources in the field. These variables include the size of the oviposition sites (e.g., diameters, circumference, surface area, volume, container depth, water depth, etc.), ambient environments (temperature, relative humidity, and canopy coverage), and water characteristics (pH, conductivity, salinity, water temperature, and total dissolved solids). Methodological details can be found in Table S1 in the Appendix.

After removing eight oviposition sites with excessive missing data, we first compared each variable individually across oviposition site groups. Because our data do not follow a normal distribution, we used the Kruskal–Wallis test and post hoc pairwise Wilcoxon rank sum test in R v3.5.0 (R development core team, 2018) with *Holm* correction for multiple comparisons. We then tested the difference between habitats or between *Ae. aegypti* present and absent sites separately, regardless of the other grouping factors. We also performed a principal component analysis (PCA) to summarize all physical variables. The multivariate differences between oviposition site groups, habitats, and *Ae. aegypti* presence status were tested by multiple response permutation procedure (MRPP) with 999 permutations. The p values for multiple comparisons were adjusted using the *Holm* method. Lastly, we attempted to identify the variables that are most differentiated in each comparison by ranking variable importance using a random forest algorithm in R package *randomForest v4.6-14* (Liaw & Wiener, 2002). Random forest is a decision-tree based classification algorithm that works well with small sample size and correlated variables (Qi, 2012).

### Characterizing oviposition sites: competition and predation

Competition and predation could influence larval development and female oviposition choice (Pamplona Lde et al., 2009; Soman & Reuben, 1970; Vonesh & Blaustein, 2010; Zahiri & Rau, 1998). To consider their effects, we counted the number of individual mosquitoes (*Ae. aegypti* as well as other mosquito species) present in each oviposition site. We also noted the presence of predatory larvae, predominately *Toxorhynchites* mosquitoes, and removed them immediately if found. We first compared the number and density of *Ae. aegypti* between habitats, using only the oviposition sites where *Ae. aegypti* was present. We also carried out an additional analysis that used mosquitoes of all species (including *Ae. aegypti*) to include possible interspecific competition effects, and included oviposition sites without *Ae. aegypti*. In La Lopé, records of other mosquito species were only available for the forest, so we only compared forest sites present versus absent of *Ae. aegypti*. We used negative-binomial models to compare mosquito numbers with habitat as the predictors, and used Kruskal–Wallis tests and *post hoc* pairwise Wilcoxon rank sum tests to compare mosquito density. In addition, we analyzed the frequency of finding predators in different oviposition site groups or habitats with chi-squared tests.

### Characterizing oviposition sites: microbial density

We examined the microbial profile in a subset of oviposition sites, inspired by previous studies showing that the microbiome, particularly bacteria, affect *Ae. aegypti* oviposition (Arbaoui & Chua, 2014; Ponnusamy et al., 2015). We collected 15 mL (in La Lopé) or 50 mL (in Rabai) water samples from each field oviposition site using sterile pipets and conical tubes (Thermo Scientific, USA). This procedure was performed before measuring physical characteristics to avoid contamination. We kept the water samples in a cooler with ice packs in the field until returning to the field station. To measure microbial density, we added an aliquot of each water sample to formaldehyde solution (Millipore Sigma, USA) with a final concentration of 1% - 3% formaldehyde and kept it in 4 °C. After returning to Yale University, we stained the formaldehyde preserves with DAPI (4’,6-diamidino-2-phenylindole, final concentration 5 ug/mL, Thermo Scientific, USA), and counted the microbial cells using hemocytometers (DHC-N01, INCYTO, Korea) under a widefield fluorescence microscope (Leica DMi8, Leica, German) Densities were log-transformed before statistical analysis. We then compared the microbial density among oviposition site groups in La Lopé with the Kruskal–Wallis test and post hoc pairwise Wilcoxon rank sum tests. The distribution of data in Rabai samples did not violate parametric test assumptions, so we performed the comparisons using analysis of variance (ANOVA) and *post hoc* Tukey tests.

### Characterizing oviposition sites: bacterial community composition

In addition to the overall density, we performed 16s-rRNA gene amplicon sequencing to explore the bacterial community composition in most oviposition sites (Table 1), inspired by previous studies suggested different bacteria between habitats (Dickson et al., 2017). The details of sample processing and sequencing library preparation are described in the Appendix. In short, we collected cells from the water samples by centrifuge or filtering, extracted DNA, and amplified the 16s-rRNA gene V4 region using primers reported in Kozich et al. (2013). The primers label each sample with a unique combination of index sequences. The PCR products were cleaned and mixed with equal quantity and sequenced on Illumina MiSeq (Illumina, USA) at the Yale Center for Genome Analysis. We also included commercial mock communitis of bacteria in our library. The composition of these mock communities are known, which allows validation of the sequencing accuracy. Amplicon sequencing for La Lopé and Rabai were conducted separately.

We demultiplexed the sequencing reads using USEARCH v10.0.240 (Edgar, 2010) and followed the pipeline of DADA2 (v1.8.0) (Callahan et al., 2016) to determine the bacterial community composition. DADA2 estimates sequencing errors and infers the exact sequence variants (i.e., amplicon sequence variants, or ASVs), which are analog to the conventional operational taxonomic unit (OTU). We summarized the frequency of each ASV in every water sample, and blasted the ASVs to the Ribosomal Database Project (RDP) 16s-rRNA gene reference database (RDP trainset 16 and RDP species assignment 16) (Cole et al., 2014) for taxonomic assignment. We then agglomerated ASVs into higher taxonomic levels for further analysis.

Using the DADA2 outputs and the R package *phyloseq* (McMurdie & Holmes, 2013), we first calculated the alpha diversity of the bacteria community in each oviposition site indicated by the Shannon index (Shannon, 1948), using the raw read counts of all samples. We then compared the index across oviposition site groups, habitats, and between *Ae. aegypti* present and *Ae. aegypti* absent sites. The community compositions were summarized by non-metric multidimensional scaling (NMDS) with the Bray-Curtis distance matrix. Similar to PCA, NMDS analysis summarizes multivariate data (each bacterial taxa as one variable), but is more appropriate for bacterial composition data (Ramette, 2007). Before NMDS analysis, we first removed samples with fewer than 5000 reads to avoid low-quality samples, and we thinned each sample proportionally to the lowest read depth of all samples to remove the impact of uneven sequencing depth between samples. Bacterial communities may show different assembly patterns at different taxonomic levels (Goldford et al., 2018). Therefore, we calculated the Shannon index and performed NMDS at four taxonomic levels: ASV, Species, Genus, and Family. To provide more information on the detailed compositions of the bacterial communities, we also demonstrated the major bacterial groups at the Family level. Lastly, we used R package *DESeq2* to identify families that are most differentiated between habitats (Love, Huber, & Anders, 2014).

To estimate the temporal stability of the bacterial communities, for five oviposition sites in each habitat, we collected water samples more than once. The average number of days between two consecutive collections ranges from 3 to 21, with an average of 8.4 days in La Lopé and 17 days in Rabai. All temporal samples were sequenced, but only the first-day samples were included in analyses described above. We performed a separate NMDS analysis to examine variation between temporal samples.

### Characterizing oviposition sites: chemical volatiles in Rabai, Kenya

Chemical volatiles released from an oviposition site could act as olfactory cues for mosquitoes during oviposition site selection (Afify & Galizia, 2015), yet the volatile profiles of natural oviposition sites have rarely been examined. We attempted to describe the volatile profile in oviposition sites in Rabai, Kenya (we did not collect chemical data in La Lopé due to financial constraints). In brief, we collected water samples from a subset of oviposition sites (Table 1) and extracted the volatiles into an absorbent with a steady airflow. The captured volatiles were examined by Gas Chromatography–Mass Spectrometry (GC-MS) at Yale West Campus Analytical Core. We then identified and quantified each compound using the GC-MS results. The technical details of volatile extraction and GC-MS were described in the Appendix. Due to the sparsity of many compounds in the final dataset, we did not perform statistical analysis across oviposition site types or habitats, but instead summarized the compound concentrations using a heatmap.

### Field oviposition choice experiments

We conducted field oviposition experiments in both La Lopé and Rabai. We placed artificial and natural containers at forest sites and village sites and left them for use by wild mosquitoes. Bamboo segments were used for the natural containers since they are similar to tree holes in size and shape and have been commonly used by African researchers to collect forest mosquitoes (Kemp & Jupp, 1991). The artificial containers used in La Lopé included tires, plastic bottles, plastic bags, bricks, and metal cans (see the insets in Figure 7 for a representation of these experimental containers). These containers are frequently found in the villages. We paired five bamboo with the five artificial containers to form a group of ten containers. We then placed these groups in four forest locations and four peridomestic locations. All containers were set up empty and filled by rainwater naturally. We retrieved all containers after roughly two weeks, collected larvae and pupae from them, and reared all mosquitoes to adults to count the number of *Ae. aegypti*. Because of the low yield in these experimental containers, within each habitats, we combined mosquitoes from all bamboos or all artificial containers, respectively. This resulted in a single count of *Ae. aegypti* from each types of container in habitat. We used a chi-squared test to examine whether habitat influences the distribution of *Ae. aegypti* in the bamboo versus artificial containers.

In Rabai, we followed similar procedures but used plastic buckets and earthenware pots as the artificial containers. Each container group thus consisted of two artificial containers and two bamboo fragments. Another difference is that instead of placing the container group in peridomestic as in La Lopé, we left them in domestic habitats (indoor), after receiving verbal permission from homeowners. We set up ten container groups in the Kaya Bomu forest and ten in Bengo village (Figure 1B). Tap water was added to the containers on the first day, as rains were not frequent enough and could not reach indoor containers. The experiment lasted for 7-10 days. In the end, containers were flooded to hatch all eggs, and we reared larvae and pupae in the field. We then counted the number of *Ae. aegypti* present in the bamboo or either type of artificial container. Instead of combining mosquito counts as described above for La Lopé experiments, we kept data from the ten containers groups (i.e. ten replicates) within each habitat separate. Each of the ten container groups thus represents a replicate in the choice experiment. After removing groups that produced no *Ae. aegypti*, we applied a beta-binomial model to address the effect of habitat on the distribution of eggs between bamboo and artificial containers. The beta-binomial model was implemented in the R package *glmmTMB* (Brooks et al., 2017).

Although we chose containers as similar as possible to natural oviposition sites in both habitats, it is critical to examine how the environmental conditions of these experimental containers reflect the natural conditions. Therefore, we collected water samples from them at the end of the experiments and applied the 16s-rRNA gene amplicon sequencing and downstream NMDS analysis to examine the bacterial community in these experimental containers.

### Laboratory oviposition assays

In an attempt to disentangle the effects of different environmental variables on oviposition choice, we performed laboratory oviposition assays in a common-garden setup. The goal was to examine whether forest and village *Ae. aegypti* have different oviposition preferences towards a subset of environmental variables that differed between forest and village oviposition sites.

We established a forest colony and a peridomestic colony from La Lopé using *Ae. aegypti* collected from natural breeding sites, supplemented with oviposition traps and human landing capture (approved by the National Research Ethics Committee of Gabon under the protocol 0031/2014/SG/CNE). In Rabai, we created six independent village colonies from the four villages (four domestic colonies and two peridomestic colonies) and four forest colonies from the the Kaya Bomu forest. We blood-fed the mosquitoes in the field and brought the eggs back (i.e., the second generation) to our lab at Yale University and the McBride lab at Princeton University. The detailed information of the mosquito colonies and the protocol for rearing these colonies are in the Appendix. The two peridomestic colonies correspond to K63 and K65 in Rose et al. (2020) and the Rabai forest colonies correspond to K66 and K67. All laboratory oviposition assays were performed in the insectary at Yale University. We used the fourth to the sixth generation of mosquitoes in these assays. For simplicity, we will refer to the peridomestic colony in La Lopé and the domestic colonies in Rabai as village colonies.

In the first series of assays, we used two-choice tests. In each cage, five gravid females were allowed to choose from two oviposition cups with different conditions (see Appendix). Using this assay, we compared the oviposition preference of forest versus village colonies from La Lopé and Rabai towards several environmental variables.

We first tested a pair of Rabai forest and domestic colony (K66 vs. Kwa Bendegwa village) for their preference for water samples directly collected from forest and domestic oviposition sites in Rabai. The rest of the oviposition assays focused on specific environmental variables. We determined the target variables from the list of variables that showed a significant difference between forest and village oviposition sites in the field. The decision was also constrained by space, resources, and our experimental setup. For example, we were unable to test variables related to container size and height as well as ambient temperature and humidity, as they require much larger space than the capacity of our insectary. As a result, we tested pH, shading, larval density, and a combined effect of pH, conductivity, and shading in the K66 versus Kwa Bendegwa colony pair. Additionally, we conducted oviposition assays examining bacterial community composition in all La Lopé and Rabai colonies (Table S2 in the Appendix).

Conditions we examined in the assays replicated the median conditions of forest and village oviposition in nature (details of each assays are described in the Appendix). Specifically, in the experiments testing the preference for bacterial compositions, we create forest and village type of bacterial community by inoculating water samples collected from natural forest and village oviposition sites in nutritionally rich Lysogeny broth (LB). After growing the two bacterial cultures overnight, they were diluted to the same cell density and used as the two choices in the behavioral assays. Although the bacterial communities in these LB cultures likely vary from the actual bacterial communities in natural oviposition sites, they should still contain some representative bacterial taxa from each habitat.

We counted the number of eggs in oviposition cups at the end of each assay. Cages with fewer than ten eggs in total were removed from further analysis. We first calculated the oviposition activity index (OAI) (Kramer & Mulla, 1979) for each cage:

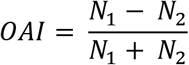

where *N_1_* and *N_2_* are the number of eggs deposited in the two cups, respectively. *OAI* ranges from −1 to 1, which represents a complete preference for the second choice to a complete preference for the first choice. We performed beta-binomial models in the R package *glmmTMB* (Brooks et al., 2017) to examine whether colonies differ in their oviposition preference, using the two egg counts in each cage as the dependent variable (Rose et al., 2020). We added the batch/trial IDs as random effects if the experiments testing a condition spanned more than one trial. The statistical significance of colony or habitat effects were determined by comparing the full model with a null model that excludes colony or habitat information (Table S9). We then extracted mean OAI with a 95% confidence interval from the model using the R package *emmeans* (Lenth, Singmann, & Love, 2018; Rose et al., 2020).

In addition to the above two-choice assays, in the last series of laboratory assays, we tested the oviposition preference of all mosquito colonies to five bacterial densities. This is inspired by the large variation in bacterial density among filed oviposition sites (more than two orders of magnitude) and that previous laboratory experiments with *Ae. aegypti* found density-dependent ovipositional responses to bacteria (Ponnusamy et al., 2015; Ponnusamy, Wesson, Arellano, Schal, & Apperson, 2010). We used a similar experimental design as the two-choise assays described above but provided each cage of gravid female mosquitoes five cups instead of two. The cups contained bacterial cultures at densities ranging from zero to nearly the maximal bacterial density in field oviposition sites. The bacterial culture was generated from an even mixture of forest and domestic water samples (Table S2 in the Appendix). We counted the numbers of eggs laid in the five cups and fitted a negative-binomial model using the R package *lme4* (Bates, Mächler, Bolker, & Walker, 2014) to detect significant differences between colonies and between the habitat type of the colonies. A full model with an interactive term of colony/habitat with bacterial densities was compared to a null model excluding this interactive term (Table S9). Because the number of eggs in the five cups from the same cage were not independent from each other, we added the ID of cages as a random effect to control for this data structure. Lastly, we used the *emmeans* package to estimate the expected number of eggs in each bacterial density with 95% confidence intervals.

## Results

### Characterizing oviposition sites: physical characteristics

PCA analysis summarizing the 11 physical variables in La Lopé showed that the four oviposition site groups (two habitats × *Ae. aegypti* present/absent) overlap extensively in the space described by the first two principal components, which together account for 38% of the total variance (Figure 2a). However, forest and peridomestic village sites appeared to differ slightly. In support of that, MRPP tests found a significant multivariate difference among the four groups (p = 0.019) and between habitats when including both *Ae. aegypti* present and absent sites (p = 0.001). Sites with *Ae. aegypti* did not differ significantly between habitats (p = 0.316), possibly due to the small sample size (only five samples in the forest *Ae. aegypti* present category). Sites with and without *Ae. aegypti* did not differ significantly (all sites regardless of habitats: p = 0.311, only forest sites: p = 1, only peridomestic sites: p = 1).

**Figure 2.**
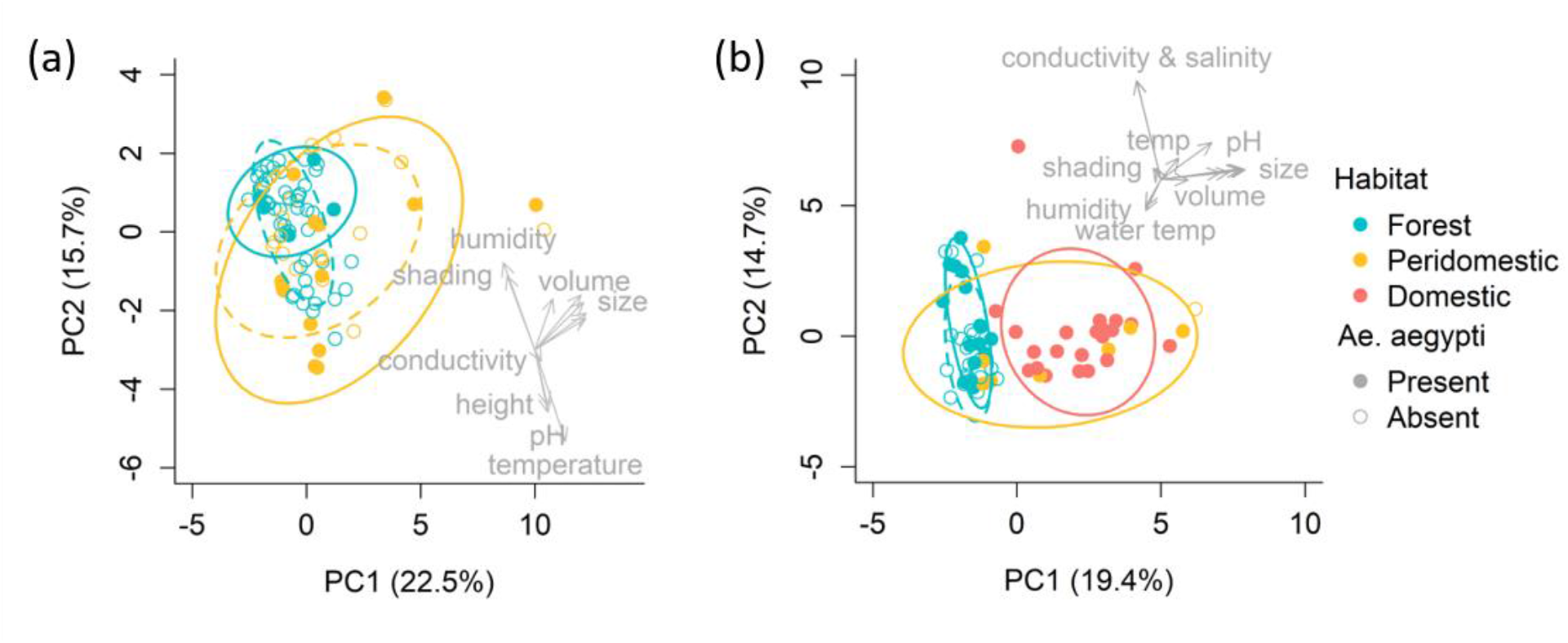
Principal component analysis (PCA) of all physical variables in (a) La Lopé and (b) Rabai. The first two PCs are shown, and the variance explained by each PC was indicated in the axis label. Each point represents a single oviposition site. Colors and point shapes indicate habitat and whether *Ae. aegypti* were found in the sites, respectively. An eclipse was drawn for each oviposition site group with a 75% confidence level. The colors of the eclipses represent habitat types and match the colors of the points. The solid and dashed eclipses correspond to *Ae. aegypti* present and absent sites. The original variables were overlaid on the PC1-PC2 plate with major variables labeled.

Examining the rotations of the original physical variables onto the first two principal component axes suggested two main groups of variables. The first one includes ambient temperature and humidity, shading (i.e., canopy coverage), container opening height, and water pH, which axis roughly corresponds to differentiation between forest and peridomestic oviposition sites. For these variables, we found a significant difference between habitats and between the four oviposition site groups (Figure S2, Table S3 and S4). They also ranks highly in their variable importance measures (Figure S4a). These observations support the idea that variables in this group differentiate habitats. The second group mainly represents container size and water volume, with the latter differing significantly between forest and peridomestic oviposition sites (Table S3). *Ae. aegypti* present and absent sites have similar conditions for all variables except the height of the container opening (Figure S2, Table S3 and S4). Container height is the main factor differentiating *Ae. aegypti* present vs. absent sites, especially within the peridomestic sites (Figure S3a).

In Rabai, similarly as in La Lopé, forest and peridomestic sites was modestly but significantly different in their physical characteristics (Figure 2b, PCA summarizing 16 physical variables). In addition to these two habitat types, we also measured indoor ‘domestic’ sites. The PCA results suggest a strong differentiation between forest and domestic oviposition sites (Figure 2b). Sites with and without *Ae. aegypti* in the forest did not show much difference. Consistent with PCA, MRPP found significant multivariate differences in most comparisons, except between forest sites with *Ae. aegypti* present vs. absent (p = 0.157) and between domestic vs. peridomestic sites (p = 0.192).

Forest and domestic oviposition sites were separated primarily along the first PC, which is explained by container size (e.g., diameter, circumference, etc.), water volume, and water pH (Figure 2b). Single variable comparisons confirmed that these variables are indeed different between forest and domestic/peridomestic oviposition sites (Figure S4, Table S5 and S6). On the other hand, comparisons of forest *Ae. aegypti* present vs. absent sites as well as between domestic and peridomestic sites found very few significant differences (Table S5 and S6). Canopy coverage, a measure of shading, also showed a strong difference across oviposition site groups and between habitats (Figure S4, Table S5 and S6). We expected this difference as domestic oviposition sites are always under roof, and peridomestic containers are mostly exposed, while forest tree holes are partially shaded by the canopy. Lastly, variables with significant differences between oviposition site groups or between habitats also generally have high ranks in the output of the Random Forests analysis of the corresponding comparisons (Figure S3b).

### Characterizing oviposition sites: competition and predation

The density of *Ae. aegypti* was similar between forest and peridomestic oviposition sites in La Lopé (Figure 3a, Table S4). Many oviposition sites in both habitats produced only one *Ae. aegypti* (Figure S5a). Although peridomestic sites contained significantly more *Ae. aegypti* than forest sites (Figure S5a), they also had larger volumes (Figure S2f). Mosquitoes other than *Ae. aegypti* were recorded only in the La Lopé forest. Oviposition sites with and without *Ae. aegypti* in the forest did not show a significant difference in the total number or density of all mosquitoes (Figure S5b and S5c, Table S4). Analysis of predation found that the presence of predatory *Toxorhynchites* larvae did not differ among oviposition site groups, between habitats, or between *Ae. aegypti* present and absent sites (p > 0.05 in all chi-squared tests).

**Figure 3.**
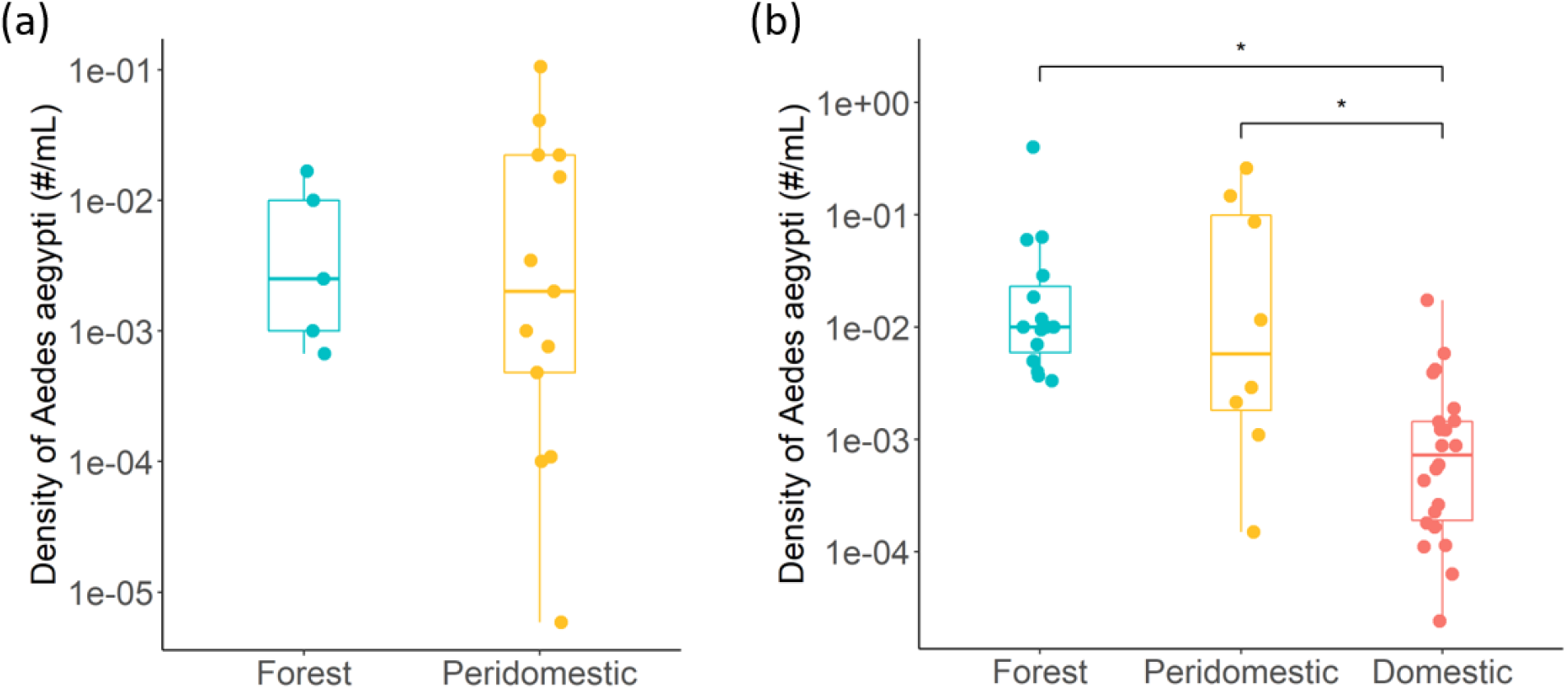
Comparison of *Ae. aegypti* density between habitats in (a) La Lopé and (b) Rabai. Only oviposition sites present with *Ae. aegypti* were included. Each point represents a single oviposition site. The color and shape are as in Figure 2. The boxplots show the minimum, 25% quartile, median, 75% quartile, and maximum. Differences between habitats were tested using pairwise Wilcoxon rank sum test with *Holm* multiple comparison corrections (*: p < 0.05, Table S4-S6).

In Rabai, *Ae. aegypti* density was significantly lower in domestic oviposition sites (Figure 3b, Table S6) in comparison with the other two habitats. The density difference between forest and domestic containers was mainly driven by the difference in water volume (Figure S4e). In contrast, the difference between peridomestic and domestic sites are due to the higher number of mosquitoes found in peridomestic sites (Figure S6a). When including other mosquito species, comparisons of mosquito numbers and densities between oviposition sites groups reached the same conclusion (Figure S6b and S6c, Table S5 and S6).

### Characterizing oviposition sites: microbial density

Microbial densities do not show significant differences between oviposition site groups, habitats, or *Ae. aegypti* present and absent sites in La Lopé (Figure 4a, Table S3 and S4). In Rabai, we found significantly lower microbial density in domestic oviposition sites than forest or peridomestic oviposition sites (Figure 4b, Table S5 and S6). Microbial densities were similar in forest and peridomestic oviposition sites. Lastly, *Ae. aegypti* present and absent sites have comparable levels of microbial density (Figure 4b, Table S5 and S6).

**Figure 4.**
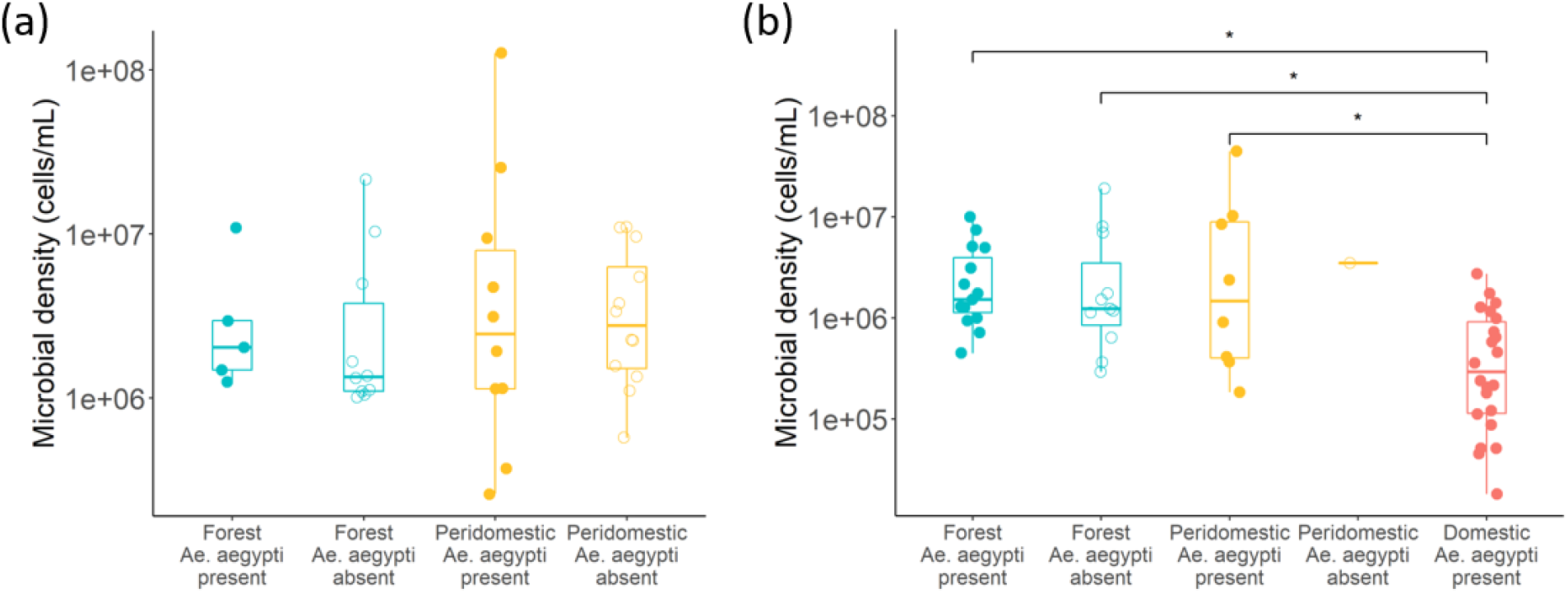
Comparison of microbial density between oviposition site groups in (a) La Lopé and (b) Rabai. Each point represents an oviposition site. The color and shape are as in Figure 2. Differences between groups were tested using pairwise Wilcoxon rank sum test with *Holm* multiple comparison correction (*: p < 0.05, Table S3-S6).

### Characterizing oviposition sites: bacterial community composition

The median depth of the amplicon sequencing was 17,420 reads per La Lopé samples and 56,478 reads per Rabai samples. Negative controls yielded 0 - 976 reads (median: 33) and 0 - 11 ASVs (median: 4) per sample, which suggested minimal contamination from the sampling and library preparation procedures. We also reconstructed the mock communities relatively well using the sequencing results: we found 18-23 ASVs from mock communities containing 20 bacterial taxa and nine ASVs from another two mock communities that contain eight bacterial colonies (see the Appendix for more information on the mock communities).

In La Lopé, alpha diversity of the bacterial communities varies considerably. The Shannon index differs significantly across the four oviposition site groups at the Species and Genus level (Table S3). The pairwise comparison did not find any significant difference between any pairs of oviposition site groups at any taxonomic level (Table S4). Still, visually the peridomestic groups have higher Shannon indexes (Figure S7). This observation was reflected in the comparisons between habitats regardless of *Ae. aegypti* presence, as it suggested a significantly higher alpha diversity in peridomestic oviposition sites at the species, genus, and family level (Table S3). *Ae. aegypti* present and absent sites have similar alpha diversity at all taxonomic levels (Table S3). In Rabai samples, we did not find significant differences in the Shannon index across oviposition site groups or between habitats (Figure S8, Table S5 and S6). *Ae. aegypti* present sites have lower diversity than *Ae. aegypti* absent sites, but only when we included oviposition sites from all habitats (Table S5).

NMDS analysis suggested that forest and village (including peridomestic and domestic) oviposition sites had a very different bacterial community in both La Lopé and Rabai at the ASV level. Peridomestic sites in Rabai clustered with domestic sites (Figure 5). The forest-village divergence was less evident at higher taxonomic levels for the La Lopé oviposition sites, especially at the Family level (Figure S9). Rabai samples, on the other hand, retained the substantial difference between forest and village oviposition sites at all four taxonomic levels (Figure S10). In all NMDS analysis, oviposition sites present and absent with *Ae. aegypti* within each habitat always overlap extensively, which suggested that they likely have similar bacterial community composition (Figure 5, S9, and S10).

**Figure 5.**
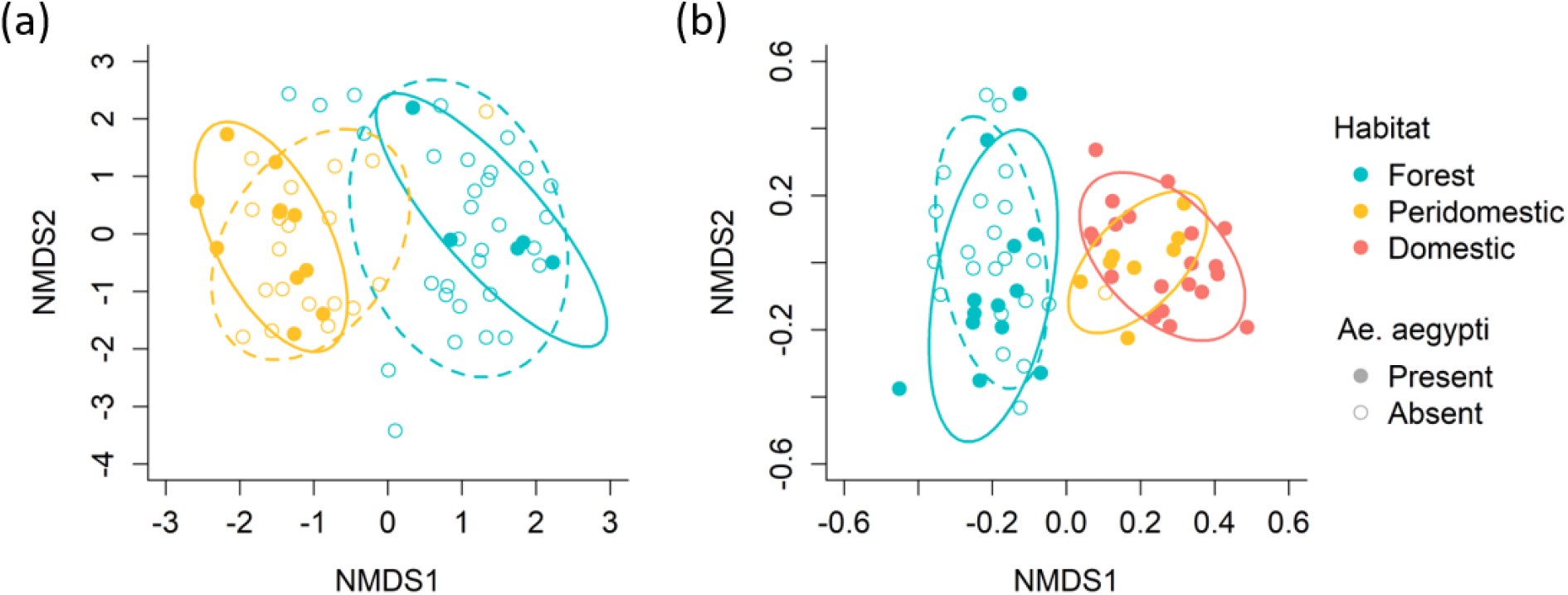
NMDS analysis of bacterial community compositions in oviposition sites in (a) La Lopé and (b) Rabai. The analysis was performed with the amplicon sequencing results at the sequencing variants (ASVs) level. Each point represents an oviposition site. The color and shape of points, as well as the ellipses, are the same as in Figure 2.

When examining the most abundant bacterial families across different oviposition site groups, we observed considerable variation among samples (Figure S11). Most oviposition sites contained representatives of multiple families with no clear dominance. Among the top ten families in La Lopé samples, *Microbacteriaceae*, *Flavobacteriaceae*, and *Burkholderiaceae* showed higher abundance in forest oviposition sites, while *Oxalobacteraceae* and *Sphingobacteriaceae* are more abundant in peridomestic sites (Figure S11a, Table S7). In Rabai oviposition sites, *Moraxellaceae* has an apparent dominance in domestic oviposition sites, but its abundance is not significantly different between habitats. *DESeq2* found a significantly higher abundance of *Enterobacteriaceae, Xanthomonadaceae*, *Pseudomonadaceae*, and *Planococcaceae* in forest oviposition sites than domestic and peridomestic sites (Figure S11b, Table S8). A full list of bacterial families that showed differential abundance between habitats are in Table S7 and S8 in the Appendix.

Lastly, NMDS analysis at the ASV level found that temporal samples collected from the same oviposition site do vary in their bacterial community, but remain in the same cluster defined by habitats (Figure S12). That is, temporal samples from forest cluster with the rest of forest oviposition sites instead of sites from other habitats and vice versa. This result suggests that the strong divergence in bacterial communities between habitat are likely temporally stable.

### Characterizing oviposition sites: chemical volatiles in Rabai, Kenya

The volatile profiles of a subset of oviposition sites in Rabai were summarized in Figure 6. After filtering, 31 oviposition sites remained in the final dataset. There was substantial variation in the chemical composition of samples, both within habitats and across habitats. GC-MS analysis identified a total of 29 chemical compounds. The majority of them were shared across different habitats, but we found a few chemicals that were unique to either forest or domestic habitat.

**Figure 6.**
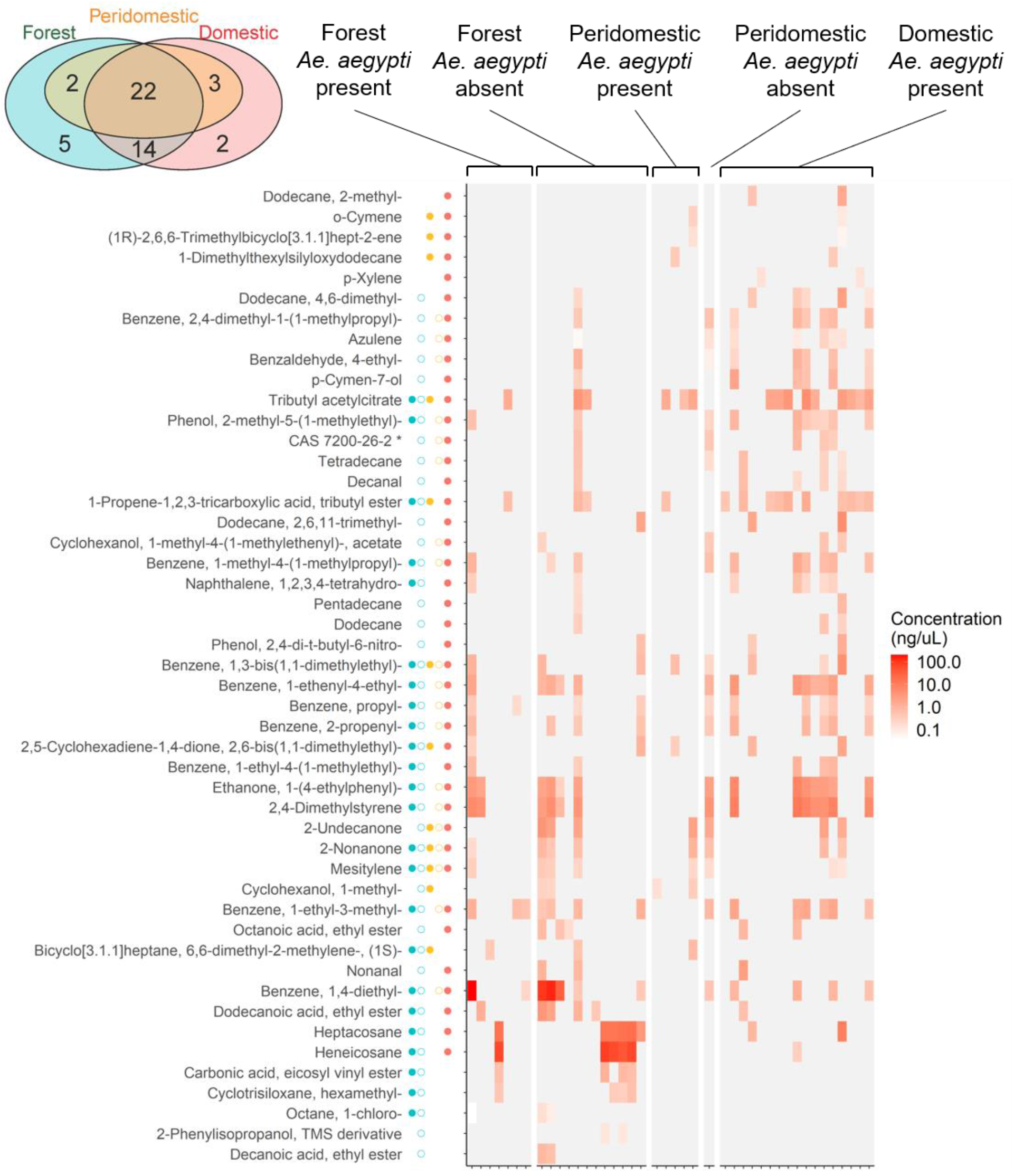
Chemical profile of the volatile samples collected from Rabai oviposition sites. Each row represents a compound, and each column represents an oviposition site. The five columns of points between the compound names and the heatmap summarize whether the compounds were present in each of the five oviposition site groups. The color and shape of points are the same as in Figure 2. The color of each cell in the heatmap quantifies the concentration on a log scale. Gray cells indicate that the compound was not found in the oviposition sites according to the GC-MS results. The inset Venn diagram shows the total numbers of compounds unique in each habitat or shared between different habitats.

### Field oviposition choice experiments

Experimental containers in the forest and the village produced in total 61 and 95 *Ae. aegypti*, respectively, in La Lopé. The majority of the *Ae. aegypti* were from bamboo in the forest and artificial containers in the village (Figure 7a). This habitat-associated bias in *Ae. aegypti* production between the two types of containers was statistically significant (chi-square test: χ^2^ = 52.1, df = 1, p < 0.001). In Rabai, we collected all but one *Ae. aegypti* from artificial containers in the village (Figure 7b). *Ae. aegypti* were also more abundant in artificial containers than in bamboo in the forest, which is the opposite of the finding in La Lopé. However, the beta-binomial model still found a significant effect of habitat (Figure 7c, AIC of full model: 54.3, AIC of null model: 48.9, model comparison: χ^2^ = 7.38, df = 1, p = 0.007).

**Figure 7.**
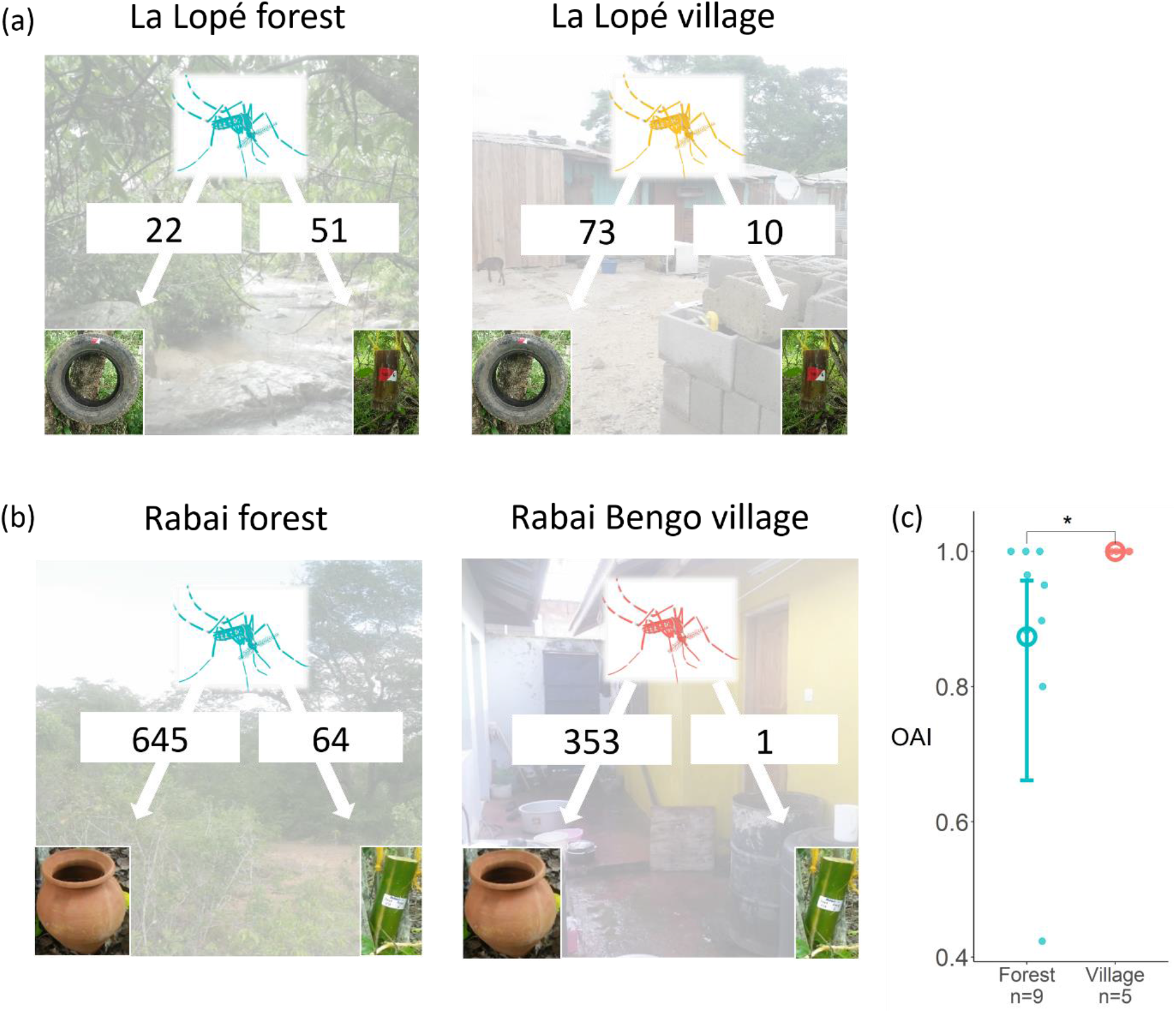
Field oviposition choice experiments in (a) La Lopé and (b) Rabai. Artificial containers and bamboo segments (inset photos as examples) were placed in both the forest and the villages. The numbers in (a) and (b) are the total numbers of *Ae. aegypti* produced by the two types of containers in the two habitats. In Rabai, the ten replicates of container pairs in each habitat were examined separately. An OAI was calculated for each container group that had no fewer than ten *Ae. aegypti*, as shown by points in (c). Larger OAI implies more *Ae. aegypti* from artificial containers. The hollow circles and error bars show the mean OAI and a 95% confidence interval estimated by a beta-binomial model with habitats as the predictor. The model was significantly better than a null model, which suggested a significant difference between habitats (*: p < 0.05).

When examining the bacterial community composition of these experimental containers, NMDS analysis found that regardless of the container type and the habitats where they were located, all experimental containers clustered with natural village (peridomestic and domestic) oviposition sites (Figure S13).

### Laboratory oviposition assays

The results of the laboratory oviposition assays are shown in Figure 8 and Figure S14. Based on the OAI confidence interval estimated by the beta-binomial models, we found three significant preferences among all experiments: Rabai Kwa Bendegwa village colony preferred forest water samples over village water samples, and forest mosquito larval density over village larval density; La Lopé forest colony preferred the bacterial culture started with peridomestic water samples over that started with forest water samples. However, there is significant within-colony variation in most experiments. When comparing between colonies or between the habitat types of the colonies, the beta-binomial models did not find any significant difference in any assays (Table S9). Lastly, we applied a negative-binomial model to analyze the results of oviposition assays testing bacterial densities (Figure 9). Neither colonies nor the habitats of the colonies have a significant effect on the mosquito’s preference for the five bacterial densities (Table S9). La Lopé village colonies showed a weak preference for lower bacterial densities, but the trend was not statistically significant (ANOVA for the effect of oviposition choices: F = 1.56, df = 4, p = 0.200).

**Figure 8.**
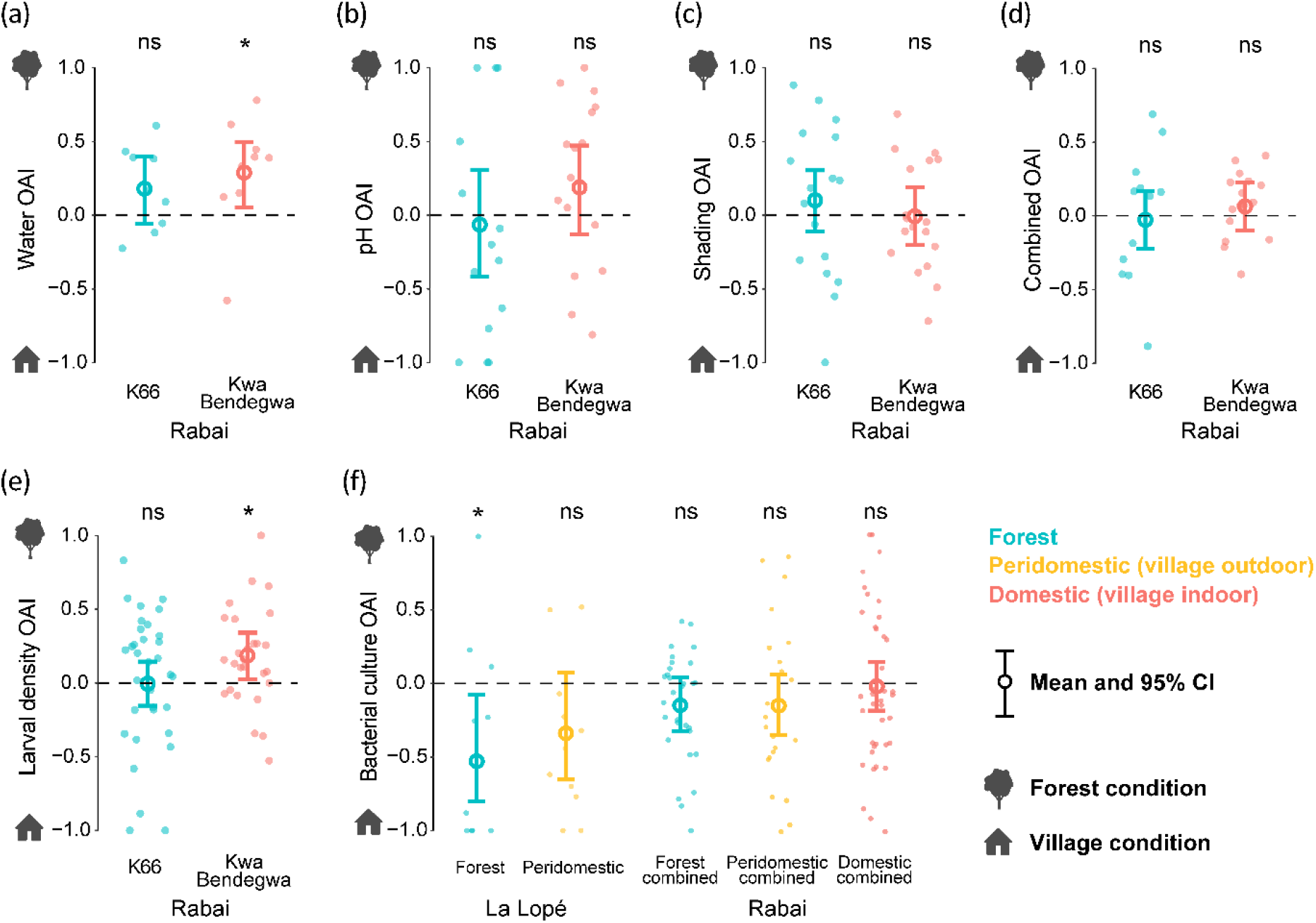
Two-choice laboratory oviposition assays testing preference for field-collected waters, pH, shading, a combination of water pH, salinity and shading, *Ae. aegypti* larval density, and bacterial culture. Colony-wise results are shown in Figure S14 in the Appendix. The details of the two choices in each assay were described in Table S2 in the Appendix. Higher OIA implies a preference for the forest condition. Each point represents the OAI of one cage with five gravid females. The mean and 95% confidence interval (CI) were estimated by beta-binomial models. The asterisks and ‘ns’ above each colony indicates whether the 95% CI excludes zero. No sigficant differences were found between habitats or between colonies in any experiments.

**Figure 9.**
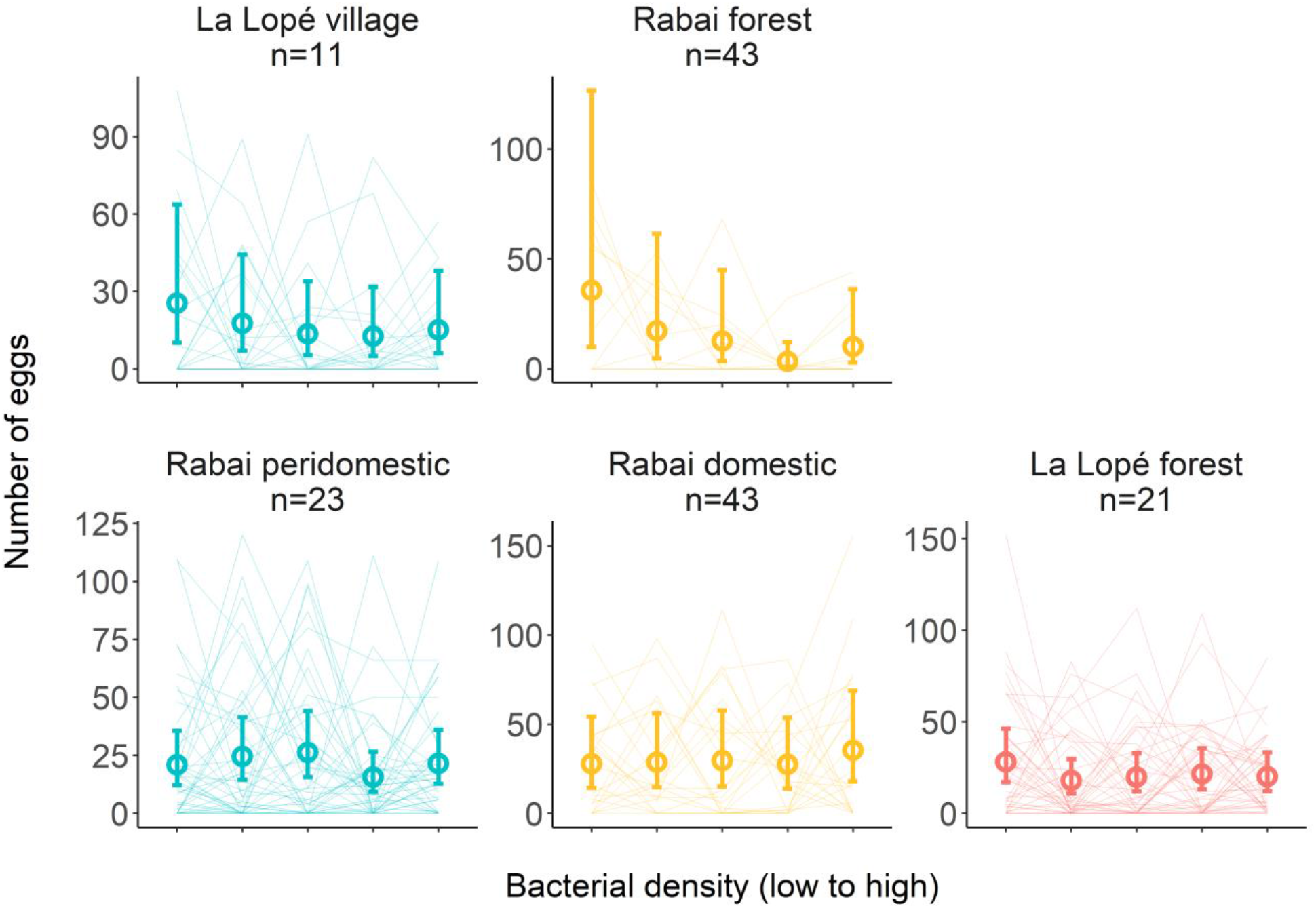
Five-choice laboratory oviposition assays testing preference for bacterial density. Five cups were provided in each cage with increasing bacterial density at 0, 2×10^5^, 1×10^6^, 5×10^6^, 2.5×10^7^ cells/mL (details in Table S2), which correspond to the five columns (left to right) in each panel. Each line connects the five egg counts in one cage. Colors represent the habitats from where the colonies came. Multiple colonies from the same habitat in Rabai were combined in this figure. Colony-wise results are shown in Figure S15 in the Appendix. A negative-binomial model was used to fit the results of each oviposition assay. The model estimates the mean number of eggs in each bacterial density and a 95% confidence interval, which are shown by the open circles and the error bars, respectively.

## Discussion

In this study, we found that oviposition sites in different habitats tend to have different physical properties and bacterial community composition, in both La Lopé and Gabon (Figure 2 and 5). Outdoor peridomestic sites have moderately different physical characteristics from forest sites in both locations, while the forest-domestic comparison unique to Rabai reveals a even stronger differentiation (Figure 2). The bacterial composition in forest oviposition sites is consistently very distinct from the other two village habitats (Figure 5). Unique to the Rabai system, we also found significantly lower larval and bacterial densities in domestic oviposition sites (Figure 3b and 4b), as well as some differences in the chemical profiles between forest and domestic oviposition sites (Figure 6). These results support our hypothesis that *Ae. aegypti* living in their ancestral forest habitats and invaded anthropogenic habitats are using oviposition sites with different average properties.

Within each habitat, oviposition sites with *Ae. aegypti* present or absent at the time of collection share similar environmental conditions. We found some significant differences between *Ae. aegypti* present and absent sites, but only when we combined sites across all habitats. These results can be partially explained by the uneven distribution of *Ae. aegypti* present vs. absent sites across habitats. For example, in Rabai, all but one *Ae. aegypti* absent sites were in the forest, so the comparison between all *Ae. aegypti* absent and present sites were largely confounded by the contrast between forest and domestic sites. Another caveat for this comparison between *Ae. aegypti* present vs. absent sites lies in the difficulty of confirming that the absence of *Ae. aegypti* in any site was due to active avoidance. However, if *Ae. aegypti* actively choose only a subset of oviposition sites with specific environments, we would expect to see a tighter clustering of sites with *Ae. aegypti* present than absent, which is not the case in our data (Figure 2 and 5). Therefore, it is likely that wild *Ae. aegypti* do not have a strong preference when choosing oviposition sites within their native habitats and could use most available sites.

Many environmental characteristics of natural oviposition sites, such as the bacterial communities, predation risk, and competition are likely dynamic and vary depending on weather, season, and stochastic events (e.g., leaf litter falling into a site). In our data, we did find variation in bacterial communities in temporal samples from a few oviposition sites. However, these temporal differences did not exceed the scale of within-habitat variation (Figure S12), which suggested that the main difference we found between forest and peridomestic/domestic oviposition sites were likely stable over time. We expect a similar or even higher level of temporal stability in most physical characteristics of oviposition sites as they are more intrinsic to the containers (e.g., container size) or their locations (e.g., shading). Unfortunately, all our sampling was conducted during the rainy season, and the largest interval between two temporal samples was 21 days, which is not enough to evaluate seasonal variabilities. In dry seasons, it is generally easier to collect *Ae. aegypti* by oviptraps in the field (personal observations), which may suggest that either the mosquitoes are less selective when there are fewer natural oviposition sites available, or the mosquitoes have an altered preference in dry seasons. Recently studies suggested that seasonality may play an important role in driving the domestication of *Ae. aegypti* (Powell & Tabachnick, 2013; Rose et al., 2020). Therefore, future studies are needed to examine the seasonal change of oviposition site conditions, in order to provide a full picture of the ecological backdrop for mosquito oviposition.

Only a few studies so far has characterized the environmental conditions of *Ae. aegypti* natural oviposition sites. Dickson et al. (2017) described the bacterail community composition in field oviposition sites in La Lopé, Gabon and found a strong differentiation between habitats, which is echoed in our study. Yet that study did not examined other environmental conditions such as the physical variables. Another study compared several environmental conditions between tree holes and tires in Hattiesburg, MS, USA, and found consistent differences between them (Yee et al., 2012). However, *Ae. aegypti* were not present in most containers in that study. Therefore, the environmental data reported in the current study added useful information to our understanding of the ecology of *Ae. aegypti* oviposition. Admittedly, our data may not cover the full temporal variations in the field, and the complex field conditions limited the accuracy of some measures. For instance, accurate volume and surface area estimates were challenging for some irregularly shaped sites. However, we hope this initial quantification of natural oviposition sites could provide useful information for generating hypotheses regarding the evolution of *Ae. aegypti* oviposition.

One hypothesis we wanted to test in this study is that environmental differentiation between forest and village (peridomestic and domestic) oviposition sites leads to divergent oviposition preference in the mosquitoes. The results of the field oviposition assay did suggest some behavioral differences between forest and village *Ae. aegypti* in both La Lopé and Rabai (Figure 7). However, these results need to be interpreted with caution. Because we counted *Ae. aegypti* after they developed into adults instead of at the egg stage, the number in each experimental container could be affected by factors other than oviposition preference, such as egg hatching rate and larval survival etc. In addition, although bamboo segments are similar to tree holes in size and shape, they possessed bacterial communities that resembled the artificial containers (Figure S13). Therefore, results of the field oviposition experiments might reflect a behavioral difference, if it truly exists, that does not fully correspond to the between-habitat difference of natural oviposition sites. It is unclear what are the exact mechanisms of this differential production of *Ae. aegypti* from bamboo and artificial containers in different habitats, which could be of interest for future studies. An interesting possibility is that choice of bamboo versus artificial containers was affected by their apparency in each habitat (Harrington et al., 2008; Strauss, Cacho, Schwartz, Schwartz, & Burns, 2015). For example, domestic habitats may present less visual obstacles and make the aritifical container stand out more. This potential visual effect may be of interest for future studies.

Nevertheless, the results of field experiments provided strong evidence that *Ae. aegypti* in the forest habitat readily accept artificial containers. They might even prefer these containers, as we collected more *Ae. aegypti* from these artificial containers placed in the forest (22 in La Lopé and 645 in Rabai) than from tree holes or rock pools (9 in La Lopé and 156 in Rabai). These results imply that *Ae. aegypti* may be predisposed to use artificial containers for oviposition. Oviposition choices have been suggested to have a strong impact on the movement of *Ae. aegypti* (Reiter, 2007). Previous studies also proposed that females turning to human stored water for oviposition during dry seasons may be a key driver for the human specialization of *Ae. aegypti* inside Africa (Brown et al., 2014; Powell et al., 2018; Powell & Tabachnick, 2013; Rose et al., 2020). As suggested by our field experiment results, this crucial ovipositional transition might happen relatively easily and frequently. Consistent with this hypothesis, a population genetic study using *Ae. aegypti* collected in La Lopé and Rabai found very little evidence of genetic differentiation between habitats, which indicates that mosquitoes could move between habitats freely (Xia et al. submitted). On the other hand, this extensive connectivity in the local scale between habitats may hinder any phenotypic divergence from evolving, consistent with the lack of oviposition differences in the lab (Figure 8). In a more regional scale where gene flow is less frequent, there may be differences between mosquitoes from different habitats, as found for host odor preference by Rose et al. (2020).

In line with this “predisposal” hypothesis, it is possible that *Ae. aegypti* from La Lopé and Rabai are not very selective in their oviposition choices in general. We found considerable OAI variation within each colony despite the well-controlled rearing and experimental procedures. Only a few trials found any significant preference for any choices. Yet in these assays, the direction of preference was opposite our prediction (e.g., one domestic colony from Rabai showed a preference for forest water samples over domestic water samples; Figure 8a). Lastly, we did not find a significant difference between forest and village mosquito colonies in any assays (Figure 8 and 9). However, these results of the laboratory oviposition assays need to be interpreted with caution. For example, we cannot rule out the possibility that that we simply lacked the power to detect the preference. Yet, our sample sizes are comparable to many previous studies that used a similar experimental design and found significant oviposition preference (Afify, Horlacher, Roller, & Galizia, 2014; Allan & Kline, 1995; Ganesan, Mendki, Suryanarayana, Prakash, & Malhotra, 2006; Melo et al., 2020). It is also possible that the choices we tested are not of a magnitude detectable by female *Ae. aegypti*. However, these choices were informed by the characteristics of natural oviposition sites, and therefore should be ecologically relevant for the mosquitoes. We are currently testing some more extreme conditions (e.g., complete shading vs. complete exposure) using the same groups of La Lopé and Rabai colonies, which will be summarized in a future report.

Another strong possibility is that mosquitoes use multiple cues simultaneously in choosing oviposition sites, as previous studies found a broad spectrum of factors influencing *Ae. aegypti* oviposition (Afify & Galizia, 2015; Arbaoui & Chua, 2014; Day, 2016; Harrington et al., 2008; Leahy et al., 1978; Wong, Stoddard, Astete, Morrison, & Scott, 2011). Because most of our choice assays focused on a single variable, it is premature to reach a definitive conclusion of no behavioral difference. Future experiments testing more combinations of environmental factors are needed to gain a deeper understanding of the potential synergistic effects of the environments on driving oviposition evolution in *Ae. aegypti formosus*.

In summary, this study confirmed a strong environmental difference between forest and village oviposition sites in both Gabon (La Lopé) and Kenya (Rabai). Our ecological divergence hypothesis suggested that *Ae. aegypti* in different habitats may evolve divergent oviposition preferences corresponding to these environmental differences. However, direct behavioral data from this study was insufficient to support this hypothesis. The similar environmental conditions between *Ae. aegypti* present vs. absent sites in the field also suggested no strong selectivity within habitats. Considering all the findings, it is possible that *Ae. aegypti* in La Lopé and Rabai behave as generalists when choosing oviposition sites. If this is the case, the initial transition between habitats may not require significant changes in oviposition behavior. After occupying different habitats, mosquitoes may start to evolve some minor behavioral differences, but likely not strong enough to discriminate against oviposition sites from the other habitats and impede gene flow at this small geographic scale. This speculation is consistent with the documentation of multiple independent invasions of domestic habitats in Africa in recent years (Kotsakiozi, Evans, et al., 2018; Powell & Tabachnick, 2013), including the latest cases of La Lopé and Rabai (Xia et al., submitted). Being an ovipositional generalist benefits *Ae. aegypti* as they are capable of utilizing a large variety of containers (Petersen, 1977; Simard, Nchoutpouen, Toto, & Fontenille, 2005), and thus quickly respond to environmental changes such as the drying of tree holes as well as within-container competition. In the forest, most oviposition sites we surveyed contained multiple species. A previous study in Kenya found a positive association between *Ae. aegypti* and a few other *Aedes* species in tree holes (Lounibos, 1981), which could lead to resource competition. It is possible that this competition and possibly predation, in combination with the flexibility of *Ae. aegypti* oviposition choices, drove the mosquito to exploit artificial containers. This raises the question of why most other mosquito species do not exploit domestic habitats. What makes *Ae. aegypti* so special?

*Ae. aegypti* are known to spread risks during oviposition by a conservative bet-hedging strategy (Starrfelt & Kokko, 2012), namely ‘skip oviposition’: A gravid female distributes her eggs across multiple containers to prevent losing all eggs due to the destruction of any single oviposition site (Colton, Chadee, & Severson, 2003; Swan, Lounibos, & Nishimura, 2018). If *Ae. aegypti* can accept a large variety of oviposition choices, they could further spread the risk. It would be interesting to examine whether the large inter-individual variation we observed in oviposition choice assays are heritable and consistent across the lifetime of individual mosquitoes.

This study examined *Ae. aegypti* in forests and rural villages in Africa, where the domestication of this epidemiologically important species likely first occurred (Powell et al., 2018; Powell & Tabachnick, 2013). Outside of Africa, *Ae. aegypti* are closely associated with human communities and use almost exclusively artificial containers for oviposition, except in the Caribbean and Argentina (Chadee et al., 1998; Mangudo et al., 2015). Studies from the 1970s and continuing through 2014 found a human-specialized strain of *Ae. aegypti* reintroduced to Rabai from America or Asia (Brown et al., 2011; McBride et al., 2013; Tabachnick et al., 1979; Tabachnick & Powell, 1978; Trpis & Hausermann, 1978), which have likely gone extinct before our study in Rabai in 2017 (Xia et al. submitted). Comparing this re-introduced strain with the local sylvatic *Ae. aegypti* back then revealed significant behavioral differences, including their oviposition preference (Leahy et al., 1978; Petersen, 1977; Trpis & Hausermann, 1975). These pieces of evidence suggested that *Ae. aegypti* outside of Africa have behaviorally specialized to the domestic oviposition sites. When did the ovipositional adaptation happen, if *Ae. aegypti* remain largely generalists during the initial invasion inside Africa? A few recent studies suggested that human specialization may happen somewhere in West Africa, such as Sahel or Angola (Crawford et al., 2017; Powell et al., 2018; Rose et al., 2020). This specialization may not always accompany the use of domestic habitats inside Africa, but may play a key role for the spread of this species to the rest of the world. More studies examining the intial domestication process inside Africa and the later human specialization are necessary for providing a more comprehensive understanding of the evolutionary history of *Ae. aegypti*.

## Supporting information

Appendix that includes method details, supplementary tables and figures

## Data Accessibility Statement

The datasets that describe the basic information, physical characteristics, larval density, predator presence, microbial density, and chemical profile of oviposition sites in La Lopé and Rabai are archived in Dryad: doi:10.5061/dryad.7m0cfxprg (La Lopé) and doi:10.5061/dryad.3tx95×6cz (Rabai), repestively. The 16s-rRNA gene amplicon sequencing data was deposited in the NCBI SRA database with ID SUB7716639 (La Lopé samples) and SUB7719551 (Rabai samples).

## Author contributions

SX and JRP designed and conceptualized the study. SX, DA, RS, JL, CSM, NHR, and JRP coordinated the fieldwork. SX, DA, and JL conducted field sampling of oviposition sites and field oviposition experiments. SX and HD designed the volatile collection in Rabai and performed the GC-MS analysis. SX performed the lab work to generate the data on microbial density and bacterial community composition. SX, CSM, and NHR established mosquito colonie, and SX performed the laboratory oviposition assays. SX wrote the first draft of the manuscript. JRP provided funding, coordinated the entire study, and interpreted results with SX. All authors provided critical feedback on the manuscript.

## Acknowledgments

We appreciate the collaboration and the support from Institut de Recherche pour le Développement (IRD) and the research Unit ESV-GAB at the Centre International de Rrecherches médicales de Franceville (CIRMF) in Gabon, and Kenya Medical Research Institute (KEMRI) in Kenya during the fieldwork. We are grateful to all the field assistants and scientists in the field, especially Nil Rahola and Marc F. Ngangue in La Lopé and Rotich Gilbert in Rabai. In addition, we thank Andrew Goodman and his lab for providing primers for bacterial amplicon sequencing and helpful guidance in library preparation. We also thank Nanxi Lu for the instructions on bioinformatic analysis of the sequencing results. We received a lot of technical support and training from the Yale Center for Genome Analysis (YCGA) on Illumina sequencing, from the West Campus Analytic Core on GC-MS, and from the Yale West Campus Imaging Core on fluorescent microscopes and we are grateful for all the support. The design of the laboratory experiments benefited greatly from the helpful discussions with Luciano Cosme, Ryan Joseph, Lisa Baik, and Noah Rose. We appreciate all the useful discussions, suggestions, and feedback from Gisella Caccone, Tom Chiodo, Benjamin Evans, Stephen Gaughran, Andrea Gloria-Soria, Evelyn Jensen, Panagiota Kotsakiozi, Joshua Miller, Evlyn Pless, Maud Quinzin, Norah Saarman, Samuel Snow, and John Soghigian. We also want to thank the McBride lab at Princeton University for valuable feedback and discussions. Lastly, we are very grateful for the advice and guidance from Stephen Stearns, John Carlson, and Alvaro Sanchez.

This work was supported by NIH RO1 AI101112 to JRP and YIBS Small Grants Program, Doctoral Dissertation Improvement Awards to SX.

