## Appendix that includes method details, supplementary tables and figures for "Oviposition of the mosquito *Aedes aegypti* in forest and domestic habitats in Africa"

### Method details

*16s-rRNA gene amplicon sequencing of water samples from natural oviposition sites*

In La Lopé, after aliquoting a subset of water samples into formaldehyde solution, we kept the water in -20 °C in the field until returning to CIRMF, Franceville, Gabon. We centrifuged the samples after thawing using a Backman TJ-6 centrifuge (Beckman Coulter, USA) at 6000 rpm (maximum speed) for 30 minutes to collect microbial cells. The liquid was removed immediately after centrifuge, and we extracted DNA using the QIAGEN Blood and Tissue kit (QIAGEN, USA) following the manufactural protocol. The DNA was stored at -20 °C until brought back to the lab at Yale University. In Rabai, due to lack of access to centrifuge, we instead used a filtering approach to collect microbiome from water samples. Specifically, upon bringing the water samples back to the field station, we pushed around 50 mL water of each sample using a sterile syringe through a Millipore Sterivex filter unit (SVGPL10RC, EMD Millipore, USA) with 0.22 µm pore size to collect all microbial cells. We then sealed the filtering unit and frozen them at -20 °C until bringing them back to Yale University. Under aseptic conditions, we pealed the members from the filter unites and extracted DNA from them using the DNeasy PowerWater Kit (QIAGEN, USA) according to the manufactural protocol.

We followed the same protocol as in Kozich et al. (2013) to prepare two sequencing libraries for samples collected in La Lopé and Rabai, respectively. The protocol amplified the V4 region of the bacterial 16s-rRNA gene with duel indexes, which allows us to multiplex multiple samples in one sequencing. The primers were prepared in Dr. Andrew Goodman's lab at Yale University as Gülden et al. (2017). PCR was conducted using the Q5 High-Fidelity DNA Polymerase (New England Biolabs Inc., USA) with a similar PCR cycle in Kozich et al. (2013). The amplification products were cleaned using SPRI beads (AMPure XP, Beckman Coulter, USA), and we determined the DNA concentrations by Qubit (Qubit Fluorometric and dsDNA HS Assay Kit, Thermal Fisher, USA). We then mixed the samples with equal quantity. To evaluate the sequencing and analysis performance, we added four positive controls into the mixed sequencing library, including two samples of genomic DNA from Microbial Mock Community B (HM-276D and HM-277D, BEI Resources, NIAID, NIH as part of the Human Microbiome Project) (Nelson, Morrison, Benjamino, Grim, & Graf, 2014), one genomic DNA from ZymoBIOMICS (D6305) and one mixed community cells from ZymoBIOMICS (D6300). The final library was then examined on Bioanalyzer (Thermo Fisher, USA) to confirm the amplicon size and sent to the Yale Center for Genome Analysis for sequencing using Illumina MiSeq (Illumina, USA).

*Characterizing the volatile chemical profiles of oviposition sites in Rabai, Kenya*

We collected 8-15 mL water samples from a subset of oviposition sites (Table 1) in glass vials previously washed and baked at 400 °C. An air pump (Casella Apex Pro, UK) was used to generate an airflow at 0.2 L/min to extract the volatiles from the water in the vials for 24 hours. The chemicals in the airflow leaving the vials were captured using Volatile Collection Trap (Volatile Collection Trap LLC, USA) with PoraPak-Q and later eluted in 200 µL Hexane (Sigma-Aldrich, USA). 1-Bromoheptane (B67570, Sigma-Aldrich, USA) were mixed in the Hexane as an internal standard with a concentration of 100 ng/µL. We started volatile extraction in the evening of each collection day, which is less than 12 hours from the water collection. An empty vial was used every day as a positive control to characterize the background chemical profile in the airflow. The solution after elusion was kept at -20 °C until shipped back to Yale. We analyzed the chemical profile using gas chromatography-mass spectrometry (GC-MS, Agilent 7890A/5975C, Agilent Technologies, Inc. USA) at Yale West Campus Analytical Core.

We analyzed the GC-MS results using MSD ChemStation F.01.03.2357 (Agilent Technologies, Inc. USA). We first identified any compounds that exist only in the samples or have a substantially higher quantity in the samples compared to the positive control. The compounds were then identified using the National Institute of Standards and Technology (NIST) reference library (v 2.2, Scientific Instrument

Services, USA). We quantified the concentrations of each compound using the area underneath the corresponding peak(s) and used the 1-Bromoheptane peak to translate area to absolute concentration. We removed compounds found in only one oviposition sites which could result from identification errors. We also excluded compounds that do not exist in natue (http://www.thegoodscentscompany.com), which suggested contamination or misidentification. After filtering the compounds, 11 oviposition sites without any of the remaining chemical compounds were also removed.

*Mosquito colonies*

*Ae. aegypti* collected from natural oviposition sites were kept alive in the field to establish colonies. In La Lopé, due to the low number of *Ae. aegypti* collected, we pooled all individuals from each habitat to establish a single forest colony and a single village colony. We also distributed several bamboo traps and performed human landing catches in La Lopé forest to supplement the larval collection. In Rabai, all colony-forming *Ae. aegypti* came from natural breeding sites. We established two forest colonies (one from the deep area in the forest and another from the edge of the forest adjacent to the Chang'ombe village) and four domestic colonies collected from the domestic oviposition sites in the four villages, respectively. In both La Lopé and Rabai, we blood-fed the females in the field multiple times, which is necessary for egg development and collected the eggs (the second generation) on seed germination papers (SD7606, Anchor Paper Company, USA). After producing eggs, *Ae. aegypti* were preserved in ethanol in the field for genetic analysis.

The eggs of the two La Lopé colonies and the four Rabai domestic colonies were brought back to Yale at the end of the fieldwork. The two Rabai forest colonies were first brought back to the McBride Lab at Princeton University and maintained as described in Rose et al. (2020) with code name K66 and K67. A copy of the third generation of these two colonies was later sent to Yale while the original copy was continued at Princeton University. We kept the mosquitoes in the insectary with 27 ˚C constant temperature, 50%-70% relative humidity, and a 12h/12h light/dark cycle. Eggs were hatched with deionized water supplied with fish food (TetraMarine Saltwater Granules, Tetra, German), and pupae were transferred into insect rearing cages (BugDorm-1) that are roughly 30 x 30 x30 cm. We provided 10% sugar water to adults constantly and fed them multiple times with sheep blood (DSB050, Hemostat, USA) at least five days after they emerged. Three days after feeding, we provided four cups (two black cups and two white cups) lined with seed papers to collect eggs from gravid females. The eggs were dried slowly in the insectary and kept for up to six months.

To control for the possible laboratory adaptations of these colonies to our specific insectary and rearing regime, in addition to the eight colonies in our lab (named "Powell" strains), we also acquired four more colonies from Rabai that had been reared in the McBride lab at Princeton University until the fifth generation (named "McBride" strains). They were maintained as described in Rose et al. (2020). These four colonies consist of the two Rabai forest colonies (K66 and K67) as well as two Rabai peridomestic colonies that were independently collected from the Chang'ombe village (K65) and the Mbarekani village (K63), respectively. In total, we examined two colonies from La Lopé and ten colonies from Rabai in our oviposition assays.

*Laboratory oviposition assay*

Each batch of the oviposition assays included two to four colonies, with at least one forest colony and one village colony, and we synchronized all colonies from hatching until the end of the experiment. The mosquitoes used for the laboratory oviposition choice assays were reared in the similar protocol as the main colonies described in the main texts, with a few changes to reduce between-experiment variations introduced by the rearing process. Firstly, after hatching, the first-instar larvae were transferred into new larval trays with the density of one larva per 5mL water and fed with a fixed amount of larval food. Secondly, we kept around 360 adults, including both sexes in a 17.5 cm cube cage (BugDorm-

4M1515) instead of the larger cage for colony maintenance. Lastly, we fed females with sheep blood only once about 5-8 days after more than 90% of pupae emerge into adults. Females were allowed to feed for one hour. We removed all males and females that did not appear engorged immediately after feeding.

Experiments started roughly 72 hours after blood-feeding. We used a binary-choice design except for experiments examining bacterial density (described in detail in the following paragraphs). Five gravid females were transferred into a 15 x 15 x 15 cm customized cage with messes covering both lateral sides and the top side (Figure S1). We used more than one animal per cage as preliminary trials with single females per cage rendered very low rates of response. We placed two or five black cups (one-oz plastic food container cups) containing 12 mL solution that differ in the variable of interest in the cage. The cups were lined with seed-germination papers for collecting eggs. Positions of the choices were selected randomly among the two or five cup positions (Figure S1). The experimental cages were kept in an environmental chamber (model PG031, Darwin Chambers Company, USA) with 27 ˚C, 70% humidity, 10 lux light intensity and a 12h/12h light/dark cycle, which is more accurately regulated than the insectary room. Locations of the cages in the incubator were determined randomly. We allowed the females to lay eggs for 24 hours and counted the number of eggs in each cup at the end of the experiments.

### Tables

**Table S1.** Methods to measure physical variables of mosquito oviposition sites in La Lopé, Gabon

| **Variable** | **Measurement method** | **Location*** |
|---|---|---|
| Diameter 1 | Measured as the longest diameter of the water surface. For tires lying horizontal, this variable is measured as the diameter of the outer circle. | Both |
| Diameter 2 | Measured as the longest diameter of the water surface that is perpendicular to diameter 1. For tires lying horizontal, this variable is measured as the diameter of the inner circle. | Both |
| Circumference | Calculated from diameter 1 and diameter 2 for containers with regular-shaped openings (e.g., round), or estimated from photos. | Both |
| Surface area | Calculated from diameter 1 and diameter 2 for containers with regular-shaped openings (e.g., round), or estimated from photos. | Both |
| Container depth | Measured as the distance from the bottom of the container to the lowest point of the entrance of the container | Rabai |
| Water depth | Measured as the distance from the bottom of the container to the water surface | Rabai |
| Volume | The exact volume was either calculated from surface area, and depth is the shape of the water body is roughly cylinder (e.g., bamboo traps, buckets, boxes, and cans), or estimated from the volume of water collected into the bottle when the site was emptied. When the measurement of exact volume was not possible, the volume was estimated by researchers in the field. | Both |
| Height | Measured as the distance between the lowest point of the entrance of the sites to the ground next to the site. For rock pools and other breeding sites that sit on the ground, the height is zero. | Both |
| Temperature difference | Ambient temperature was measured by HOBO UX100-011 Temperature and Relative Humidity Loggers (Onset, MA, USA). A logger was placed beside the oviposition site until the read stabilize. Another logger calibrated to the first one was placed in the field station to record the diurnal fluctuation of ambient temperature, which was then subtracted from the temperature measured by the oviposition sites to calculate the temperature difference. | Both |

| | | |
|---|---|---|
| Humidity difference | Relative humidity was measured by HOBO UX100-011 Temperature and Relative Humidity Loggers (Onset, MA, USA). A logger was placed beside the oviposition site until the read stabilize. Another logger calibrated to the first one was placed in the field station to record the diurnal fluctuation of relative humidity, which was then subtracted from the humidity measured by the oviposition sites to calculate the humidity difference. | Both |
| Canopy coverage | Measured by a spherical densiometer (Convex Model A, Forestry supply # 43887). The densiometer was held as close to the entrance of the breeding site as possible. Canopy coverage was estimated facing north, west, south, and east. The mean value of the four directions was used for future analysis. The direction was determined by a compass in the forest. | Both |
| pH[+] | **La Lopé:** Measured by a WTW-3110 pH-meter (Xylem, USA) from water samples collected in 50 mL sterile conical tubes less than 24 hours from collection. The water samples are kept in an icebox or at 4 degrees before measuring and recovered to room temperature when measuring.<br>**Rabai:** Measured by a Hach Pocket Pro+ Multi 2 Tester (Hatch, USA). | Both |
| Conductivity[+] | **La Lopé:** Measured by a WTW-3310 conductivity-meter (Xylem, USA) from water samples collected in 50 mL sterile conical tubes less than 24 hours from collection. The water samples are kept in an icebox or at 4 degrees before measuring and recovered to room temperature when measuring.<br>**Rabai:** Measured by a Hach Pocket Pro+ Multi 2 Tester (Hatch, USA). | Both |
| Salinity | Measured on-site by a Hach Pocket Pro+ Multi 2 Tester (Hatch, USA) | Rabai |
| Total dissolved solids (TDS) | Measured on-site by a Hach Pocket Pro+ Multi 2 Tester (Hatch, USA) | Rabai |
| Water temperature | Measured on-site by a Hach Pocket Pro+ Multi 2 Tester (Hatch, USA) | Rabai |

[*] This column indicates whether this variable was measured in both La Lopé and Rabai (“Both”), or only in Rabai (“Rabai”).

[+] pH and conductivity were measured slightly differently in La Lopé and Rabai.

**Table S2.** Details of laboratory oviposition assays

| **Variable** | **Methods to prepare oviposition choices** | **The choice resembling forest oviposition sites** | **The choice resembling village oviposition sites** | **Mosquito colonies** |
|---|---|---|---|---|
| Water samples collected in the field | About 10 mL water samples were collected from randomly selected 18 oviposition sites in Rabai forest and 18 in Rabai villages. The water samples were frozen in -20 ˚C. The forest and village water samples were randomly paired to create 18 pairs and used to in an oviposition assay with nine cages of forest mosquitoes and nine cages of domestic mosquitoes. | Water samples collected directly from Rabai forest oviposition sites, including sites present and absent of *Ae. aegypti*. | Water samples collected directly from Rabai domestic oviposition sites, including sites present and absent of Ae. aegypti. | Kwa Bendegwa domestic colony, Powell strain;<br><br>Rabai forest deep colony, Powell strain |
| pH | We adjusted the pH of 1x Phosphate-buffered saline (PBS) solution using hydrochloric acid (HCl) and sodium hydroxide (NaOH) to the desired value. | pH = 7.18 at the beginning of the experiment, which roughly equals the median pH of forest oviposition sites in Rabai present with Ae. aegypti (pH = 7.2). The pH at the end of the experiment is 7.1. | pH = 8.71 at the beginning of the experiment, which is slightly higher than the median pH of domestic oviposition sites in Rabai present with Ae. aegypti (pH = 8.4). The pH at the end of the experiment is 7.8. | Kwa Bendegwa domestic colony, Powell strain;<br><br>Rabai forest deep colony, Powell strain |
| Shading | We placed the experimental cage in the center of a 17.8 x 17.8 x 17.8 cm cardboard box, which allows light coming into the cage only from the top. The top side of the box was divided evenly into two halves, each covering one of the two cups in the cage. The two halves were modified to represent the shading of the forest and village oviposition sites in Rabai. To maximize the difference in shading, we placed the two oviposition cups against the opposite walls of the cage instead of 7.6 cm away as in other experiments. | The "forest" half of the top side of the cardboard box has 30 holes of 0.8 cm diameter wide, which in total counts for ~ 8% of the surface area. This condition mimics the median canopy coverage in Rabai forest oviposition sites present with *Ae. aegypti* (92%). | The "domestic" half of the top side of the cardboard box has no holes mimics the complete canopy coverage in most Rabai domestic oviposition sites present with *Ae. aegypti*. | Kwa Bendegwa domestic colony, Powell strain;<br><br>Rabai forest deep colony, Powell strain |

| | | | | |
|---|---|---|---|---|
| Combination of pH, salinity, and shading | We adjusted the pH of distilled water using hydrochloric acid (HCl) and sodium hydroxide (NaOH) and adjusted the conductivity using sodium chloride (NaCl). Different shading conditions were created, as described above. | pH = 7.11, conductivity = 901 μL, shading = 92% | pH = 8.30, conductivity = 503 μL, shading = 100% | Kwa Bendegwa domestic colony, Powell strain;<br><br>Rabai forest deep colony, Powell strain |
| Larval density | We hatched eggs of a Rabai forest strain and a Rabai village domestic strain simultaneously. The second-instar larvae of each strain were transferred to two new trays with 800 mL distilled water at different larval densities that represent the conditions of forest vs. domestic oviposition sites. We added 50 pallets of fish food per tray on the same day of larvae transferring and removed all larvae after three days. The water holding larvae of the two strains at the same larval density were mixed in equal quantity and used as the choices in the oviposition assay. | Water holding larvae for 3 days at the density of 50 larvae / 800 mL (1 larva / 16 mL). This larval density roughly matches the median larval density of all mosquito species in Rabai forest oviposition sites present with *Ae. aegypti*. | Water holding larvae for 3 days at a density of 1 larva / 800 mL. This larval density roughly matches the median larval density of Rabai domestic oviposition sites present with *Ae. aegypti*. | Kwa Bendegwa domestic colony, Powell strain;<br><br>Rabai forest deep colony, Powell strain |

| | | | | |
|---|---|---|---|---|
| Bacterial community composition * | During fieldwork, we preserved water samples from a subset of oviposition sites that were present with *Ae. aegypti* using 80% glycerol. These samples include 10 from La Lopé village, 5 from La Lopé forest, 10 from Rabai forest, and 10 from Rabai village. The glycerol preservation allows the bacteria to stay alive with minimal changes over time. We kept the preservations in -80 ˚C.<br><br>Glycerol preservations from each habitat in each location were mixed with equal quantity to create in total four mixed stocks. We inoculated the forest and village mixed stock in 10 mL Lysogeny broth (LB). The bacterial cultures were shaken at 200 rpm in 37 ˚C for 24 hours. The cell densities of the two cultures were measured by OD600 light absorption. We diluted the bacterial cultures to $1.25 \times 10^9$ cells/mL in 10 mL LB media, and then added 500 mL sterilized water. The final cell density is $2.5 \times 10^7$ cells/mL. The diluted bacterial solutions made from the forest vs. village bacterial stocks were used as the two oviposition choices. | The diluted bacterial culture generated using bacterial glycerol samples collected from forest oviposition sites present with *Ae. aegypti*. | The diluted bacterial culture generated using bacterial glycerol samples collected from village oviposition sites present with *Ae. aegypti*. | All colonies |
| Bacterial density * | We generated bacterial cultures following the same protocol as described above but inoculated the same LB media with both the forest and the village bacterial stocks. After the 24-hour growth, the bacterial culture was diluted using fresh LB to four different cell densities: $1.25 \times 10^9$, $2.5 \times 10^8$, $5 \times 10^7$, and $1 \times 10^7$ cells/mL, and then added to 500 mL sterilized water. The final cell densities are $2.5 \times 10^7$, $5 \times 10^6$, $1 \times 10^6$, and $2 \times 10^5$ cells/mL. A fifth choice is 10 mL LB in 500 mL sterilized water with no bacteria. | The median bacterial density in Rabai forest oviposition sites present with *Ae. aegypti* is $1.5 \times 10^6$ cells/mL. | The median bacterial density in Rabai forest oviposition sites present with *Ae. aegypti* is $3.0 \times 10^5$ cells/mL. | All colonies |

* We used the two La Lopé bacterial stocks in oviposition assays with the La Lopé mosquito colonies, and Rabai bacterial stocks in oviposition assays with the Rabai mosquitoes.

**Table S3.** Single variable comparisons across oviposition site groups, habitats, and *Ae. aegypti* present vs. absent sites in La Lopé

| Variable | All oviposition site groups[*] | Forest vs. Peridomestic[+] | *Ae. aegypti* absent vs. present[+] |
|---|---|---|---|
| Longest diameter | $\chi$2 = 1.54, df = 3, p = 0.673 | W = 1127, p = 0.231 | W = 619.5, p = 0.777 |
| Second diameter | $\chi$2 = 2.79, df = 3, p = 0.425 | W = 842, p = 0.257 | W = 786.5, p = 0.163 |
| Circumference | $\chi$2 = 0.52, df = 3, p = 0.915 | W = 995.5, p = 0.905 | W = 698, p = 0.617 |
| Surface area | $\chi$2 = 0.98, df = 3, p = 0.806 | W = 909.5, p = 0.563 | W = 733, p = 0.394 |
| Height of container opening | **$\chi$2 = 23.59, df = 3, p < 0.001** [#] | **W = 540.5, p < 0.001** | **W = 885.5, p = 0.003** |
| Water volume | $\chi$2 = 6.33, df = 3, p = 0.096 | **W = 727.5, p = 0.038** | W = 839, p = 0.054 |
| Temperature difference | **$\chi$2 = 15.64, df = 3, p = 0.001** | **W = 501.5, p < 0.001** | W = 739.5, p = 0.359 |
| Humidity difference | **$\chi$2 = 19.56, df = 3, p < 0.001** | **W = 1515, p < 0.001** | W = 480, p = 0.091 |
| Canopy coverage | **$\chi$2 = 10.2, df = 3, p = 0.017** | **W = 1334, p = 0.004** | W = 515, p = 0.181 |
| pH | $\chi$2 = 1.48, df = 3, p = 0.687 | W = 912.5, p = 0.58 | W = 741.5, p = 0.348 |
| Conductivity | $\chi$2 = 2.34, df = 3, p = 0.506 | W = 1089, p = 0.376 | W = 611, p = 0.713 |
| Microbial density (log transformed) | $\chi$2 = 1.33, df = 3, p = 0.722 | W = 138, p = 0.412 | W = 170, p = 0.889 |
| Shannon index at ASV level | $\chi$2 = 3.68, df = 3, p = 0.298 | W = 487, p = 0.108 | W = 508, p = 0.218 |
| Shannon index at Species level | **$\chi$2 = 8.71, df = 3, p = 0.033** | **W = 387, p = 0.005** | W = 501, p = 0.257 |
| Shannon index at Genus level | **$\chi$2 = 8.54, df = 3, p = 0.036** | **W = 390, p = 0.006** | W = 493, p = 0.307 |
| Shannon index at Family level | $\chi$2 = 7.37, df = 3, p = 0.061 | **W = 411, p = 0.012** | W = 491, p = 0.321 |

[*] Kruskal-Wallis rank sum test
[+] Wilcoxon rank sum test
[#] Statistically significant results were marked in bold

**Table S4.** Pairwise single variable comparisons between oviposition site groups in La Lopé

| Variable | Forest *Ae. aegypti* absent vs. Forest *Ae. aegypti* present[*] | Forest *Ae. aegypti* absent vs. Peridomestic *Ae. aegypti* absent[*] | Forest *Ae. aegypti* absent vs. Peridomestic *Ae. aegypti* present[*] | Forest *Ae. aegypti* present vs. Peridomestic *Ae. aegypti* absent[*] | Forest *Ae. aegypti* present vs. Peridomestic *Ae. aegypti* present[*] | Peridomestic *Ae. aegypti* absent vs. Peridomestic *Ae. aegypti* present[*] |
|---|---|---|---|---|---|---|
| Longest diameter | W = 131.5<br>p = 1 | W = 666.5<br>p = 1 | W = 352.5<br>p = 1 | W = 70<br>p = 1 | W = 38<br>p = 1 | W = 142.5<br>p = 1 |
| Second diameter | W = 112<br>p = 1 | W = 552<br>p = 1 | W = 207<br>p = 0.392 | W = 56.5<br>p = 1 | W = 26.5<br>p = 1 | W = 127<br>p = 1 |
| Circumference | W = 125<br>p = 1 | W = 610.5<br>p = 1 | W = 286<br>p = 1 | W = 69.5<br>p = 1 | W = 29.5<br>p = 1 | W = 136.5<br>p = 1 |
| Surface area | W = 119.5<br>p = 1 | W = 565.5<br>p = 1 | W = 252<br>p = 1 | W = 62<br>p = 1 | W = 30<br>p = 1 | W = 133.5<br>p = 1 |
| Height of container opening | W = 111<br>p = 1 | **W = 366.5**<br>**p = 0.005** [#] | **W = 126**<br>**p < 0.001** | W = 36<br>p = 0.802 | W = 12<br>p = 0.233 | W = 89.5<br>p = 0.164 |
| Water volume | W = 104<br>p = 1 | W = 490.5<br>p = 1 | **W = 161**<br>**p = 0.047** | W = 56.5<br>p = 1 | W = 19.5<br>p = 1 | W = 128.5<br>p = 1 |
| Temperature difference | W = 127<br>p = 1 | **W = 296.5**<br>**p = 0.005** | W = 175<br>p = 0.097 | W = 16<br>p = 0.071 | W = 14<br>p = 0.452 | W = 150.5<br>p = 1 |
| Humidity difference | W = 121<br>p = 1 | **W = 863**<br>**p = 0.004** | **W = 497**<br>**p = 0.007** | W = 96<br>p = 0.239 | **W = 59**<br>**p = 0.041** | W = 174<br>p = 1 |
| Canopy coverage | W = 98.5<br>p = 1 | W = 730<br>p = 0.4 | W = 458<br>p = 0.062 | W = 90<br>p = 0.522 | W = 56<br>p = 0.138 | W = 194.5<br>p = 1 |
| pH | W = 123.5<br>p = 1 | W = 578<br>p = 1 | W = 250.5<br>p = 1 | W = 62<br>p = 1 | W = 22<br>p = 1 | W = 122.5<br>p = 1 |
| Conductivity | W = 160<br>p = 1 | W = 680<br>p = 1 | W = 325<br>p = 1 | W = 56<br>p = 1 | W = 28<br>p = 1 | W = 136<br>p = 1 |
| Density of *Ae. aegypti* | - [+] | - [+] | - [+] | - [+] | W = 33.5<br>p = 0.961 | - [+] |

| | | | | | | |
|---|---|---|---|---|---|---|
| Density of all mosquitoes | W = 121.5<br>p = 0.723 | - [+] | - [+] | - [+] | - [+] | - [+] |
| Microbial density (log transformed) | W = 33<br>p = 1 | W = 76<br>P = 1 | W = 57<br>P = 1 | W = 35<br>P = 1 | W = 24<br>P = 1 | W = 55<br>P = 1 |
| Shannon index at ASV level | W = 78<br>p = 1 | W = 314<br>P = 1 | W = 105<br>P = 0.524 | W = 52<br>P = 1 | W = 16<br>P = 1 | W = 86<br>P = 1 |
| Shannon index at Species level | W = 92<br>p = 1 | W = 259<br>P = 0.270 | W = 81<br>P = 0.087 | W = 36<br>P = 1 | W = 11<br>P = 0.595 | W = 87<br>P = 1 |
| Shannon index at Genus level | W = 94<br>p = 1 | W = 262<br>P = 0.305 | W = 84<br>P = 0.113 | W = 34<br>P = 1 | W = 10<br>P = 0.452 | W = 88<br>P = 1 |
| Shannon index at Family level | W = 94<br>p = 1 | W = 279<br>P = 0.576 | W = 86<br>P = 0.134 | W = 35<br>P = 1 | W = 11<br>P = 0.595 | W = 89<br>P = 1 |

[*] Wilcoxon rank sum tests with multiple comparison correction using the *holm* method
[#] Statistically significant results were marked in bold
[+] Data not available for this comparison

**Table S5**. Single variable comparisons across oviposition site groups, habitats, and *Ae. aegypti* present vs. absent sites in Rabai

| Variable | All oviposition site groups[*] | Habitats[*] | Forest vs. Peridomestic[**] | Forest vs. Domestic[**] | Peridomestic vs. Domestic[**] | *Ae. aegypti* present vs. *Ae. aegypti* absent[**] |
|---|---|---|---|---|---|---|
| Longest diameter | **χ2 = 44<br>df = 3<br>p < 0.001** [#] | **χ2 = 42.74<br>df = 2<br>p < 0.001** | **W = 48.5<br>p = 0.003** | **W = 1<br>p < 0.001** | W = 82<br>p = 1 | **W = 867<br>p < 0.001** |
| Second diameter | **χ2 = 47.32<br>df = 3<br>p < 0.001** | **χ2 = 45.51<br>df = 2<br>p < 0.001** | **W = 29<br>p < 0.001** | **W = 4.5<br>p < 0.001** | W = 64.5<br>p = 0.416 | **W = 909<br>p < 0.001** |
| Circumference | **χ2 = 45.98<br>df = 3<br>p < 0.001** | **χ2 = 44.37<br>df = 2<br>p < 0.001** | **W = 37.5<br>p = 0.001** | **W = 0<br>p < 0.001** | W = 82<br>p = 1 | **W = 884.5<br>p < 0.001** |
| Surface area | **χ2 = 46.5<br>df = 3<br>p < 0.001** | **χ2 = 44.69<br>df = 2<br>p < 0.001** | **W = 34<br>p = 0.001** | **W = 1<br>p < 0.001** | W = 81<br>p = 1 | **W = 890.5<br>p < 0.001** |
| Water volume | **χ2 = 42.89<br>df = 3<br>p < 0.001** | **χ2 = 42.87<br>df = 2<br>p < 0.001** | **W = 57.5<br>p = 0.008** | **W = 0<br>p < 0.001** | W = 61<br>p = 0.31 | **W = 837.5<br>p < 0.001** |
| Container depth | **χ2 = 36.61<br>df = 3<br>p < 0.001** | **χ2 = 37.55<br>df = 2<br>p < 0.001** | **W = 85.5<br>p = 0.077** | **W = 25<br>p < 0.001** | **W = 40.5<br>p = 0.035** | **W = 780.5<br>p = 0.001** |
| Height of container openings | **χ2 = 9.53<br>df = 3<br>p = 0.023** | **χ2 = 10.14<br>df = 2<br>p = 0.006** | **W = 275<br>p = 0.008** | W = 480<br>p = 0.767 | **W = 41.5<br>p = 0.039** | W = 372.5<br>p = 0.061 |
| Water depth | χ2 = 3.66<br>df = 3<br>p = 0.301 | χ2 = 3.63<br>df = 2<br>p = 0.163 | W = 181<br>p = 1 | W = 295<br>p = 0.24 | W = 68<br>p = 0.551 | W = 563<br>p = 0.559 |

| | | | | | | |
|---|---|---|---|---|---|---|
| Temperature difference | $\chi2 = 1.77$<br>df = 3<br>p = 0.622 | $\chi2 = 2.82$<br>df = 2<br>p = 0.244 | W = 115.5<br>p = 0.485 | W = 420.5<br>p = 1 | W = 139.5<br>p = 0.244 | W = 535.5<br>p = 0.82 |
| Humidity difference | $\chi2 = 0.83$<br>df = 3<br>p = 0.842 | $\chi2 = 1.48$<br>df = 2<br>p = 0.478 | W = 201.5<br>p = 1 | W = 390.5<br>p = 1 | W = 69<br>p = 0.597 | W = 564<br>p = 0.551 |
| Canopy coverage | **$\chi2 = 48.6$**<br>**df = 3**<br>**p < 0.001** | **$\chi2 = 49.8$**<br>**df = 2**<br>**p < 0.001** | **W = 282**<br>**p = 0.004** | **W = 2.5**<br>**p < 0.001** | **W = 0**<br>**p < 0.001** | **W = 735**<br>**p = 0.004** |
| pH | **$\chi2 = 28.1$**<br>**df = 3**<br>**p < 0.001** | **$\chi2 = 27.84$**<br>**df = 2**<br>**p < 0.001** | **W = 24.5**<br>**p < 0.001** | **W = 121**<br>**p < 0.001** | W = 93.5<br>p = 1 | **W = 812.5**<br>**p < 0.001** |
| Conductivity | $\chi2 = 6.27$<br>df = 3<br>p = 0.099 | $\chi2 = 4.06$<br>df = 2<br>p = 0.131 | W = 221<br>p = 0.405 | W = 513<br>p = 0.295 | W = 87<br>p = 1 | W = 489.5<br>p = 0.721 |
| Salinity | $\chi2 = 6.34$<br>df = 3<br>p = 0.096 | $\chi2 = 3.96$<br>df = 2<br>p = 0.138 | W = 218.5<br>p = 0.461 | W = 514.5<br>p = 0.28 | W = 89<br>p = 1 | W = 495<br>p = 0.775 |
| Total dissolved solids | $\chi2 = 6.66$<br>df = 3<br>p = 0.083 | $\chi2 = 4.32$<br>df = 2<br>p = 0.115 | W = 223<br>p = 0.363 | W = 516<br>p = 0.267 | W = 87<br>p = 1 | W = 489<br>p = 0.717 |
| Water temperature | $\chi2 = 7.25$<br>df = 3<br>p = 0.064 | $\chi2 = 4.73$<br>df = 2<br>p = 0.094 | W = 203<br>p = 0.955 | W = 546<br>p = 0.089 | W = 96<br>p = 1 | **W = 313.5**<br>**p = 0.008** |
| Density of *Ae. aegypti* | - + | **$\chi2 = 20.961$**<br>**df = 2**<br>**p < 0.001** | - + | - + | - + | - + |
| Density of all mosquitoes | **$\chi2 = 37.25$**<br>**df = 3**<br>**p < 0.001** | **$\chi2 = 37.24$**<br>**df = 2**<br>**p < 0.001** | W = 228<br>p = 0.091 | **W = 794**<br>**p < 0.001** | **W = 160**<br>**p = 0.013** | **W = 755.5**<br>**p = 0.002** |

| | | | | | | |
|---|---|---|---|---|---|---|
| Microbial density (log transformed) | **F = 8.43<br>df = 3<br>p < 0.001** ^ | **F = 13.47<br>df = 2<br>p < 0.001** ^ | diff = -0.141<br>p = 0959 ^ | **diff = 1.825<br>p < 0.001** ^ | **diff = 1.966<br>p = 0.001** ^ | t = -1.914<br>p = 0.070 ^^ |
| Shannon index at ASV level | χ2 = 1.90<br>df = 3<br>p = 0.593 | χ2 = 2.610<br>df = 2<br>p = 0.271 | W = 105<br>p = 0.275 | W = 369<br>p = 1 | W = 119<br>p = 1 | W = 496<br>p = 0.787 |
| Shannon index at Species level | χ2 = 3.44<br>df = 3<br>p = 0.329 | χ2 = 1.25<br>df = 2<br>p = 0.536 | W = 160<br>p = 1 | W = 471<br>p = 0.967 | W = 119<br>p = 1 | W = 622<br>p = 0.179 |
| Shannon index at Genus level | χ2 = 4.91<br>df = 3<br>p = 0.178 | χ2 = 2.73<br>df = 2<br>p = 0.256 | W = 176<br>p = 1 | W = 509<br>p = 0.336 | W = 122<br>p = 1 | W = 650<br>p = 0.087 |
| Shannon index at Family level | χ2 = 5.69<br>df = 3<br>p = 0.128 | χ2 = 3.72<br>df = 2<br>p = 0.156 | W = 199<br>p = 1 | W = 522<br>p = 0.218 | W = 121<br>p = 1 | **W = 671<br>p = 0.046** |

[*] Kruskal-Wallis rank sum test
[**] Wilcoxon rank sum tests with multiple comparison correction using the *holm* method
[+] Data not available for this comparison
[#] Statistically significant results were marked in bold
[^] One-way Analysis of variance (ANOVA) with Tukey post hoc multiple comparisons of means
[^^] Student’s t-test

**Table S6.** Pairwise single variable comparisons between oviposition site groups in Rabai

| Variable | Forest<br>*Ae. aegypti*<br>absent<br>vs.<br>Forest<br>*Ae. aegypti*<br>present[*] | Forest<br>*Ae. aegypti*<br>absent<br>vs.<br>Peridomestic<br>*Ae. aegypti*<br>present[*] | Forest<br>*Ae. aegypti*<br>absent<br>vs.<br>Domestic<br>*Ae. aegypti*<br>present[*] | Forest<br>*Ae. aegypti*<br>present<br>vs.<br>Peridomestic<br>*Ae. aegypti*<br>present[*] | Forest<br>*Ae. aegypti*<br>present<br>vs.<br>Domestic<br>*Ae. aegypti*<br>present[*] | Peridomestic<br>*Ae. aegypti*<br>present<br>vs.<br>Domestic<br>*Ae. aegypti*<br>present[*] |
|---|---|---|---|---|---|---|
| Longest diameter | W = 99.5<br>p = 0.262 | **W = 23.5**<br>**p = 0.016** [#] | **W = 0**<br>**p < 0.001** | W = 25<br>p = 0.153 | **W = 1**<br>**p < 0.001** | W = 60<br>p = 1 |
| Second diameter | W = 82<br>p = 0.061 | **W = 10**<br>**p = 0.002** | **W = 1**<br>**p < 0.001** | W = 19<br>p = 0.051 | **W = 3.5**<br>**p < 0.001** | W = 51.5<br>p = 0.546 |
| Circumference | W = 91<br>p = 0.136 | **W = 14.5**<br>**p = 0.004** | **W = 0**<br>**p < 0.001** | W = 23<br>p = 0.11 | **W = 0**<br>**p < 0.001** | W = 60<br>p = 1 |
| Surface area | W = 85.5<br>p = 0.087 | **W = 14**<br>**p = 0.003** | **W = 0**<br>**p < 0.001** | W = 20<br>p = 0.065 | **W = 1**<br>**p < 0.001** | W = 59<br>p = 1 |
| Water volume | W = 132.5<br>p = 1 | **W = 29**<br>**p = 0.036** | **W = 0**<br>**p < 0.001** | W = 28.5<br>p = 0.271 | **W = 0**<br>**p < 0.001** | W = 47<br>p = 0.338 |
| Container depth | W = 163.5<br>p = 1 | W = 48.5<br>p = 0.403 | **W = 24.5**<br>**p < 0.001** | W = 32.5<br>p = 0.485 | **W = 0.5**<br>**p < 0.001** | W = 40.5<br>p = 0.164 |
| Height of container openings | W = 203<br>p = 1 | **W = 146**<br>**p = 0.042** | W = 311<br>p = 0.646 | W = 92.5<br>p = 0.233 | W = 169<br>p = 1 | W = 41.5<br>p = 0.184 |
| Water depth | W = 175<br>p = 1 | W = 96<br>p = 1 | W = 185.5<br>p = 1 | W = 61<br>p = 1 | W = 109.5<br>p = 0.528 | W = 61<br>p = 1 |
| Temperature difference | W = 151<br>p = 1 | W = 66<br>p = 1 | W = 237.5<br>p = 1 | W = 49.5<br>p = 1 | W = 183<br>p = 1 | W = 117.5<br>p = 1 |
| Humidity difference | W = 152<br>p = 1 | W = 96.5<br>p = 1 | W = 220.5<br>p = 1 | W = 70<br>p = 1 | W = 170<br>p = 1 | W = 69<br>p = 1 |
| Canopy coverage | W = 155.5<br>p = 1 | W = 143<br>p = 0.063 | **W = 1.5**<br>**p < 0.001** | **W = 102**<br>**p = 0.044** | **W = 1**<br>**p < 0.001** | **W = 0**<br>**p < 0.001** |

| | | | | | | |
|---|---|---|---|---|---|---|
| pH | W = 129.5<br>p = 1 | **W = 12<br>p = 0.001** | **W = 60<br>p < 0.001** | **W = 7.5<br>p = 0.005** | **W = 61<br>p = 0.008** | W = 86.5<br>p = 1 |
| Conductivity | W = 119.5<br>p = 0.983 | W = 113<br>p = 1 | W = 282<br>p = 1 | W = 90<br>p = 0.333 | W = 231<br>p = 0.25 | W = 69<br>p = 1 |
| Salinity | W = 116.5<br>p = 0.824 | W = 109<br>p = 1 | W = 283.5<br>p = 1 | W = 91.5<br>p = 0.27 | W = 231<br>p = 0.256 | W = 71<br>p = 1 |
| Total dissolved solids | W = 118<br>p = 0.902 | W = 113<br>p = 1 | W = 284<br>p = 1 | W = 92<br>p = 0.24 | W = 232<br>p = 0.231 | W = 69<br>p = 1 |
| Water temperature | W = 215.5<br>p = 0.726 | W = 118<br>p = 0.997 | W = 352<br>p = 0.06 | W = 71.5<br>p = 1 | W = 194<br>p = 1 | W = 77<br>p = 1 |
| Density of *Ae. aegypti* | - | - | - | W = 69<br>p = 1 | **W = 19<br>p < 0.001** | **W = 140<br>p = 0.040** |
| Density of all mosquitoes | W = 161.5<br>p = 0.926 | W = 114<br>p = 0.711 | **W = 467<br>p < 0.001** | W = 78<br>p = 0.711 | **W = 327<br>p < 0.001** | **W = 148<br>p = 0.015** |
| Microbial density (log-transformed) | diff = -0.139<br>p = 0.994 ^ | diff = - 0.158<br>p = 0.994 ^ | **diff = - 1.745<br>p = 0.005** ^ | diff = 0.019<br>p = 0.999 ^ | **diff = -1.884<br>p < 0.001** ^ | **diff = -1.903<br>p = 0.006** ^ |
| Shannon index at ASV level | W = 178<br>p = 1 | W = 58<br>p = 1 | W = 226<br>p = 1 | W = 43<br>p = 1 | W = 143<br>p = 1 | W = 102<br>p = 1 |
| Shannon index at Species level | W = 214<br>p = 0.809 | W = 90<br>p = 1 | W = 301<br>p = 1 | W = 42<br>p = 1 | W = 170<br>p = 1 | W = 109<br>p = 1 |
| Shannon index at Genus level | W = 212<br>p = 0.912 | W = 98<br>p = 1 | W = 326<br>p = 0.296 | W = 46<br>p = 1 | W = 183<br>p = 1 | W = 113<br>p = 1 |
| Shannon index at Family level | W = 207<br>p = 1 | W = 115<br>p = 1 | W = 336<br>p = 0.163 | W = 52<br>p = 1 | W = 186<br>p = 1 | W = 113<br>p = 1 |

[*] Wilcoxon rank sum tests with multiple comparison correction using the *holm* method

[#] Statistically significant results were marked in bold

[^] Tukey post hoc multiple comparisons of means after one-way Analysis of variance (ANOVA)

**Table S7.** Bacterial families with significantly different abundance between forest and village peridomestic oviposition sites in La Lopé

| Class: Order | Family | Frequency in forest samples | Frequency in village samples | Proportion in forest samples | Proportion in village samples | log2 fold change * | p value * |
|---|---|---|---|---|---|---|---|
| Betaproteobacteria: Unknown | Unknown | 1 | 2 | 0.015 | 0.002 | -1.592 | 0.031 |
| Flavobacteriia: Flavobacteriales | Flavobacteriaceae | 2 | 2 | 0.216 | 0 | -1.671 | 0.038 |
| Betaproteobacteria: Burkholderiales | Unknown | 11 | 0 | 0.035 | 0 | -1.950 | 0.019 |
| Unknown: Unknown | Unknown | 4 | 0 | 0.035 | 0 | -2.182 | < 0.001 |
| Betaproteobacteria: Methylophilales | Methylophilaceae | 7 | 1 | 0.017 | 0 | -2.277 | 0.043 |
| Spartobacteria: Unknown | Unknown | 7 | 0 | 0.001 | 0 | -2.335 | 0.002 |
| Armatimonadetes_gp5: Unknown | Unknown | 4 | 0 | 0.001 | 0 | -3.045 | < 0.001 |
| Betaproteobacteria: Burkholderiales | Burkholderiaceae | 27 | 4 | 0.069 | 0.027 | -3.212 | 0.003 |
| Cyanobacteria: Family_XIII | GpXIII | 1 | 0 | 0.002 | 0 | -3.261 | 0.049 |
| Gammaproteobacteria: Methylococcales | Methylococcaceae | 1 | 0 | 0.014 | 0 | -3.345 | 0.022 |
| Gammaproteobacteria: Aeromonadales | Aeromonadaceae | 9 | 0 | 0.002 | 0 | -3.389 | 0.004 |
| Chlamydiia: Chlamydiales | Parachlamydiaceae | 1 | 0 | 0.025 | 0 | -3.530 | < 0.001 |
| Deltaproteobacteria: Bdellovibrionales | Bacteriovoracaceae | 5 | 0 | 0.019 | 0 | -3.562 | 0.002 |
| Chlamydiia: Chlamydiales | Simkaniaceae | 4 | 0 | 0.007 | 0 | -3.672 | 0.004 |
| Cyanobacteria: Unknown | Unknown | 1 | 0 | 0.016 | 0 | -3.709 | 0.005 |

| | | | | | | | |
|---|---|---|---|---|---|---|---|
| Cyanobacteria: Family_IX | GpIX | 5 | 0 | 0.002 | 0 | -4.646 | 0.002 |
| Holophagae: Holophagales | Holophagaceae | 8 | 0 | 0.002 | 0 | -4.737 | < 0.001 |
| Actinobacteria: Actinomycetales | Microbacteriaceae | 14 | 1 | 0.068 | 0.001 | -4.976 | < 0.001 |
| Cyanobacteria: Family_XI | GpXI | 1 | 0 | 0.232 | 0 | -5.853 | 0.004 |
| Bacilli: Bacillales | Bacillales_Incertae_Sedis_XII | 0 | 4 | 0 | 0.048 | 23.199 | < 0.001 |
| Alphaproteobacteria: Rhodospirillales | Reyranella | 0 | 1 | 0 | 0.221 | 5.351 | < 0.001 |
| Actinobacteria: Actinomycetales | Nocardiaceae | 0 | 3 | 0 | 0.011 | 4.502 | < 0.001 |
| Verrucomicrobiae: Verrucomicrobiales | Verrucomicrobiaceae | 0 | 2 | 0 | 0.024 | 3.585 | < 0.001 |
| Oligoflexia: Oligoflexales | Oligoflexaceae | 1 | 9 | 0.001 | 0.002 | 3.459 | 0.004 |
| Acidobacteria_Gp4: Aridibacter | Unknown | 1 | 4 | 0 | 0.005 | 3.442 | 0.019 |
| Acidobacteria_Gp3: Gp3 | Unknown | 0 | 8 | 0 | 0.003 | 3.255 | < 0.001 |
| Actinobacteria: Actinomycetales | Geodermatophilaceae | 0 | 6 | 0 | 0.002 | 3.231 | 0.026 |
| Alphaproteobacteria: Rhizobiales | Xanthobacteraceae | 0 | 7 | 0 | 0.028 | 2.776 | 0.005 |
| Alphaproteobacteria: Caulobacterales | Caulobacteraceae | 10 | 20 | 0.002 | 0.007 | 2.268 | < 0.001 |
| Gammaproteobacteria: Pseudomonadales | Pseudomonadaceae | 0 | 1 | 0 | 0.085 | 2.188 | 0.037 |
| Betaproteobacteria: Burkholderiales | Oxalobacteraceae | 0 | 19 | 0 | 0.071 | 2.041 | 0.007 |
| Alphaproteobacteria: Rhizobiales | Methylobacteriaceae | 10 | 21 | 0.001 | 0.002 | 2.002 | 0.001 |
| Sphingobacteriia: Sphingobacteriales | Sphingobacteriaceae | 0 | 2 | 0 | 0.030 | 1.944 | 0.019 |

| | | | | | | | |
|---|---|---|---|---|---|---|---|
| Alphaproteobacteria: Rhodospirillales | Acetobacteraceae | 0 | 7 | 0 | 0.022 | 1.163 | 0.049 |
| Epsilonproteobacteria: Campylobacterales | Campylobacteraceae | 0 | 1 | 0 | 0.056 | -4.509 | 0.019 |

[*] Calculate by R package *DESeq2*

**Table S8.** Bacterial families with significantly different abundance between forest and village (domestic, peridomestic) oviposition sites in Rabai

| Class: Order | Family | Frequency in forest samples | Frequency in village samples | Proportion in forest samples | Proportion in village samples | log2 fold change * | p value * |
|---|---|---|---|---|---|---|---|
| Deltaproteobacteria: Desulfuromonadales | Geobacteraceae | 1 | 0 | 0.248 | 0 | 6.559 | < 0.001 |
| Betaproteobacteria: Rhodocyclales | Rhodocyclaceae | 2 | 0 | 0.180 | 0 | 5.855 | < 0.001 |
| Bacilli: Bacillales | Paenibacillaceae_1 | 12 | 0 | 0.003 | 0 | 5.512 | < 0.001 |
| Methanobacteria: Methanobacteriales | Methanobacteriaceae | 4 | 0 | 0.001 | 0 | 5.378 | < 0.001 |
| Deltaproteobacteria: Desulfobacterales | Desulfobulbaceae | 2 | 0 | 0.002 | 0 | 5.187 | 0.001 |
| Clostridia: Clostridiales | Peptococcaceae_1 | 1 | 0 | 0.008 | 0 | 4.987 | < 0.001 |
| Alphaproteobacteria: Rhizobiales | Methylocystaceae | 3 | 0 | 0.026 | 0 | 4.970 | < 0.001 |
| Clostridia: Unknown | Unknown | 2 | 0 | 0.001 | 0 | 4.546 | 0.001 |
| Actinobacteria: Coriobacteriales | Coriobacteriaceae | 3 | 0 | 0.002 | 0 | 4.161 | 0.019 |
| Actinobacteria: Actinomycetales | Cellulomonadaceae | 7 | 0 | 0.015 | 0 | 4.136 | 0.004 |
| Clostridia: Clostridiales | Unknown | 5 | 0 | 0 | 0 | 4.080 | < 0.001 |
| Deltaproteobacteria: Desulfovibrionales | Desulfovibrionaceae | 4 | 0 | 0 | 0 | 4.017 | 0.007 |
| Bacilli: Lactobacillales | Streptococcaceae | 7 | 3 | 0.021 | 0.003 | 3.689 | 0.044 |
| Gammaproteobacteria: Pseudomonadales | Pseudomonadaceae | 15 | 7 | 0.034 | 0.005 | 3.637 | < 0.001 |
| Clostridia: Clostridiales | Ruminococcaceae | 4 | 0 | 0.001 | 0 | 3.470 | < 0.001 |

| | | | | | | | |
|---|---|---|---|---|---|---|---|
| Clostridia: Clostridiales | Clostridiales_Incertae_Sedis_XIII | 2 | 0 | 0 | 0 | 3.401 | 0.020 |
| Clostridia: Clostridiales | Heliobacteriaceae | 1 | 0 | 0.001 | 0 | 3.248 | 0.002 |
| Bacteroidia: Bacteroidales | Unknown | 1 | 0 | 0.001 | 0 | 3.173 | 0.039 |
| Gammaproteobacteria: Enterobacteriales | Enterobacteriaceae | 10 | 2 | 0.043 | 0.001 | 3.095 | 0.003 |
| Actinobacteria: Actinomycetales | Thermomonosporaceae | 1 | 0 | 0.001 | 0 | 2.946 | 0.035 |
| Bacilli: Bacillales | Unknown | 30 | 16 | 0.008 | 0.002 | 2.717 | 0.001 |
| Actinobacteria: Actinomycetales | Streptomycetaceae | 20 | 1 | 0.007 | 0 | 2.501 | 0.002 |
| Negativicutes: Selenomonadales | Veillonellaceae | 3 | 0 | 0.001 | 0 | 2.412 | 0.008 |
| Bacilli: Bacillales | Planococcaceae | 17 | 7 | 0.020 | 0.002 | 2.347 | 0.023 |
| Gammaproteobacteria: Xanthomonadales | Xanthomonadaceae | 19 | 16 | 0.025 | 0.002 | 2.121 | 0.003 |
| Clostridia: Clostridiales | Clostridiaceae_1 | 7 | 1 | 0.002 | 0 | 1.758 | 0.020 |
| Unknown: Unknown | Unknown | 1 | 0 | 0.011 | 0 | -1.526 | 0.033 |
| Bacteroidia: Bacteroidales | Porphyromonadaceae | 0 | 2 | 0 | 0.011 | 2.979 | 0.010 |
| Planctomycetia: Planctomycetales | Planctomycetaceae | 0 | 2 | 0 | 0.010 | -1.584 | 0.030 |
| Verrucomicrobiae: Verrucomicrobiales | Verrucomicrobiaceae | 2 | 2 | 0 | 0.005 | -1.844 | 0.032 |
| Alphaproteobacteria: Rhizobiales | Methylobacteriaceae | 3 | 6 | 0 | 0.012 | -2.102 | 0.030 |
| Deltaproteobacteria: Bdellovibrionales | Bdellovibrionaceae | 0 | 2 | 0 | 0.003 | -2.132 | 0.023 |
| Actinobacteria: Actinomycetales | Micrococcaceae | 3 | 10 | 0 | 0.002 | -2.320 | 0.008 |

| | | | | | | | |
|---|---|---|---|---|---|---|---|
| Alphaproteobacteria: Rhodospirillales | Acetobacteraceae | 0 | 4 | 0 | 0.018 | -2.554 | 0.001 |
| Alphaproteobacteria: Rhodobacterales | Rhodobacteraceae | 0 | 12 | 0 | 0.027 | -2.666 | < 0.001 |
| Betaproteobacteria: Unknown | Unknown | 0 | 6 | 0 | 0.007 | -2.993 | 0.030 |
| Cytophagia: Cytophagales | Unknown | 0 | 3 | 0 | 0.001 | -3.327 | 0.023 |
| Gammaproteobacteria: Legionellales | Legionellaceae | 0 | 1 | 0 | 0.014 | -3.568 | < 0.001 |
| Cytophagia: Cytophagales | Cytophagaceae | 0 | 6 | 0 | 0.027 | -3.677 | < 0.001 |
| Unknown: Unknown | Unknown | 0 | 12 | 0 | 0.005 | -3.708 | < 0.001 |
| Flavobacteriia: Flavobacteriales | Cryomorphaceae | 0 | 3 | 0 | 0.007 | -3.833 | 0.008 |
| Chloroplast: Chloroplast | Bacillariophyta | 0 | 5 | 0 | 0.002 | -5.268 | 0.005 |
| Bacilli: Bacillales | Bacillales_Incertae_Sedis_XII | 2 | 20 | 0.001 | 0.022 | -5.770 | < 0.001 |
| Cytophagia: Cytophagales | Cyclobacteriaceae | 0 | 10 | 0 | 0.006 | -5.853 | 0.001 |
| Betaproteobacteria: Methylophilales | Methylophilaceae | 0 | 10 | 0 | 0.006 | -6.034 | < 0.001 |
| Deinococci: Deinococcales | Deinococcaceae | 0 | 18 | 0 | 0.001 | -6.269 | < 0.001 |
| Alphaproteobacteria: Rhodospirillales | Reyranella | 0 | 9 | 0 | 0.004 | -6.701 | < 0.001 |

[*] Calculate by R package *DESeq2*

**Table S9.** Results of statistical models for laboratory oviposition assays

| Variables | Colonies | Models [*] | Model comparison results |
|---|---|---|---|
| Water samples from the field | Kwa Bendegwa domestic & Rabai forest deep | Full model: egg counts in two cups ~ **Colony**<br>Null model: egg counts in two cups ~ 1 | Full model AIC: 226.69<br>Null model AIC: 225.76<br>$\chi^2$ = 1.08, df = 1, p = 0.300 |
| pH | | | Full model AIC: 162.55<br>Null model AIC: 161.07<br>$\chi^2$ = 0.52, df = 1, p = 0.472 |
| Shading | | | Full model AIC: 340.11<br>Null model AIC: 38.68<br>$\chi^2$ = 0.58, df = 1, p = 0.447 |
| Combination of pH, conductivity, and shading | | | Full model AIC: 271.42<br>Null model AIC: 269.93<br>$\chi^2$ = 0.52, df = 1, p = 0.472 |
| Larval density | Kwa Bendegwa domestic & Rabai forest deep | Full model: egg counts in two cups ~ **Colony** + (1\|Experiment ID) [+]<br>Null model: egg counts in two cups ~ 1 + (1\|Experiment ID) | Full model AIC: 566.40<br>Null model AIC: 567.39<br>$\chi^2$ = 2.98, df = 1, p = 0.084 |
| Bacterial community | La Lopé forest & La Lopé village | Full model: egg counts in two cups ~ **Colony**<br>Null model: egg counts in two cups ~ 1 | Full model AIC: 156.60<br>Null model AIC: 155.19<br>$\chi^2$ = 0.59, df = 1, p = 0.44 |
| | All Rabai colonies | Full model: egg counts in two cups ~ **Habitat** + (1\| Colony) + (1\|Experiment ID) [++]<br>Null model: egg counts in two cups ~ 1 + (1\| Colony) + (1\|Experiment ID) | Full model AIC: 816.87<br>Null model AIC: 814.46<br>$\chi^2$ = 0.55, df = 1, p = 0.451 |
| | | Full model: egg counts in two cups ~ **Colony** + (1\|Experiment ID)<br>Null model: egg counts in two cups ~ 1 + (1\|Experiment ID) | Full model AIC: 820.15<br>Null model AIC: 810.46<br>$\chi^2$ = 8.32, df = 9, p = 0.503 |
| Bacterial density | La Lopé forest & | Full model: egg count of each cup ~ **Colony x Oviposition choice** + | Full model AIC: 1039.6<br>Null model AIC: 1033.9 |

| | | | |
|---|---|---|---|
| | La Lopé village | (1\| Cage ID)<br>Null model: egg count of each cup ~<br>**Colony + Oviposition choice** +<br>(1\| Cage ID) | $\chi^2$ = 2.29, df = 4, p = 0.683 |
| | All Rabai colonies | Full model: egg count of each cup ~<br>**Habitat x Oviposition choice** +<br>(1\| Colony) + (1\|Experiment ID) + (1\| Cage ID) +++<br>Null model: egg count of each cup ~<br>**Habitat + Oviposition choice** +<br>(1\| Colony) + (1\|Experiment ID) + (1\| Cage ID) | Full model AIC: 4386<br>Null model AIC: 4374<br>$\chi^2$ = 3.98, df = 8, p = 0.858 |
| | | Full model: egg count of each cup ~<br>**Colony x Oviposition choice** +<br>(1\|Experiment ID) + (1\| Cage ID)<br>Null model: egg count of each cup ~<br>**Colony + Oviposition choice** +<br>(1\|Experiment ID) + (1\| Cage ID) | Full model AIC: 4416.8<br>Null model AIC: 4370.7<br>$\chi^2$ = 25.9, df = 36, p = 0.893 |

[*] Negative-binomial models for testing bacterial density and beta-binomial models for the rest of the oviposition assays
[+] For oviposition assays that were conducted in multiple experiment cycles, experiment ID was included as a random effect
[++] Colonies were included as random factors in statistical models examining the effects of habitats
[+++] Cage ID were included in oviposition assays to control for the paired structure of the five egg counts in each cage

**Figures**

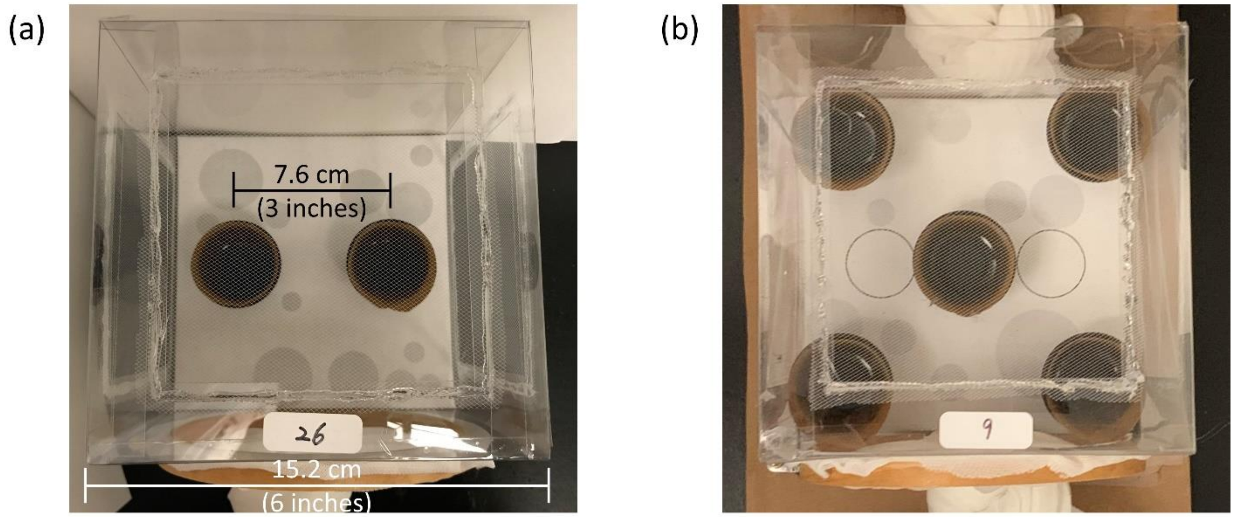


**Figure S1.** Set up of the laboratory oviposition assays with (a) two choices for testing most variables and (b) five choices for testing bacterial density. The cage was built from a 15.2 x 15.2 x 15.2 cm transparent plastic box with fine meshes covering the top and the two lateral sides. A cloth sleeve was attached to the opening on the front side (bottom in the photos) through which the mosquitoes were introduced. The cage has a white bottom with randomly generated gray circles that aims to provide visual stimuli for the mosquitoes to navigate in the cage. The black cups, each lined with a piece of 4 x 13 cm seed germination paper, were 7.6 cm away from each other in the two-choice set-up (a). In the five-choice experiments (b), the five cups located at the four corners of the cage and the center.

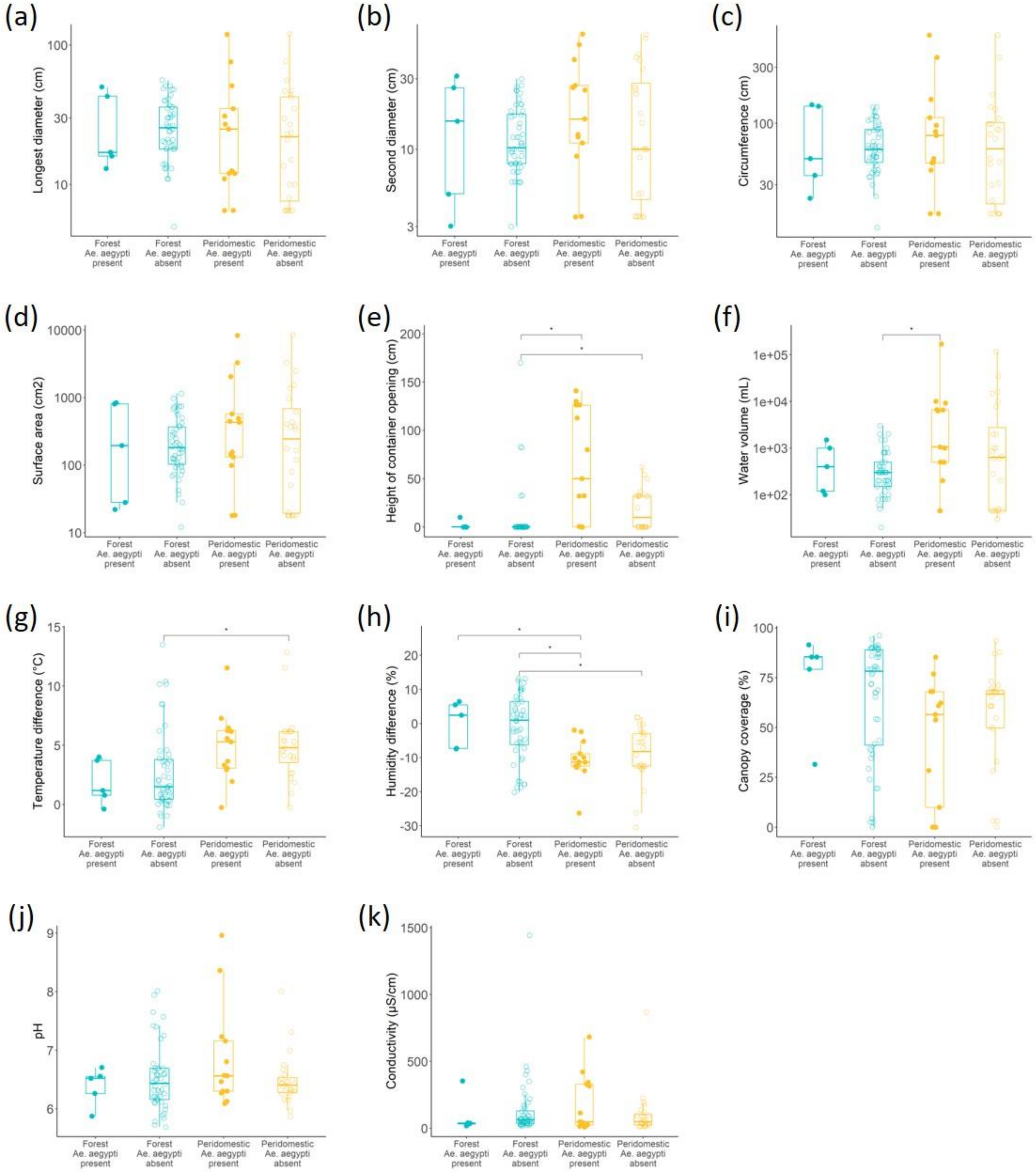


**Figure S2.** Comparison of individual physical variables between oviposition site groups in La Lopé. Each point represents a single oviposition site, and the boxplots show the minimum, 25% quartile, median, 75% quartile, and maximum of all values. The colors and shapes are as in Figure 2. Differences between groups were tested using pairwise Wilcoxon rank sum test with *Holm* multiple comparison corrections (*: $p < 0.05$, Table S3 and S4). Only significant comparisons are labeled.

(a)

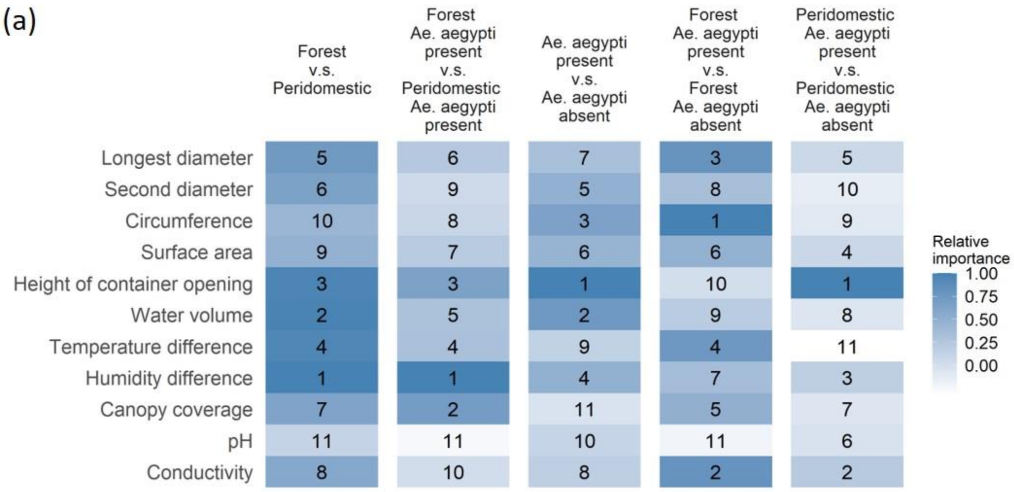

Forest v.s. Peridomestic
Forest Ae. aegypti present v.s. Peridomestic Ae. aegypti present
Ae. aegypti present v.s. Ae. aegypti absent
Forest Ae. aegypti present v.s. Forest Ae. aegypti absent
Peridomestic Ae. aegypti present v.s. Peridomestic Ae. aegypti absent
Longest diameter 5 6 7 3 5
Second diameter 6 9 5 8 10
Circumference 10 8 3 1 9
Surface area 9 7 6 6 4
Height of container opening 3 3 1 10 1
Water volume 2 5 2 9 8
Temperature difference 4 4 9 4 11
Humidity difference 1 1 4 7 3
Canopy coverage 7 2 11 5 7
pH 11 11 10 11 6
Conductivity 8 10 8 2 2
Relative importance
1.00
0.75
0.50
0.25
0.00


(b)

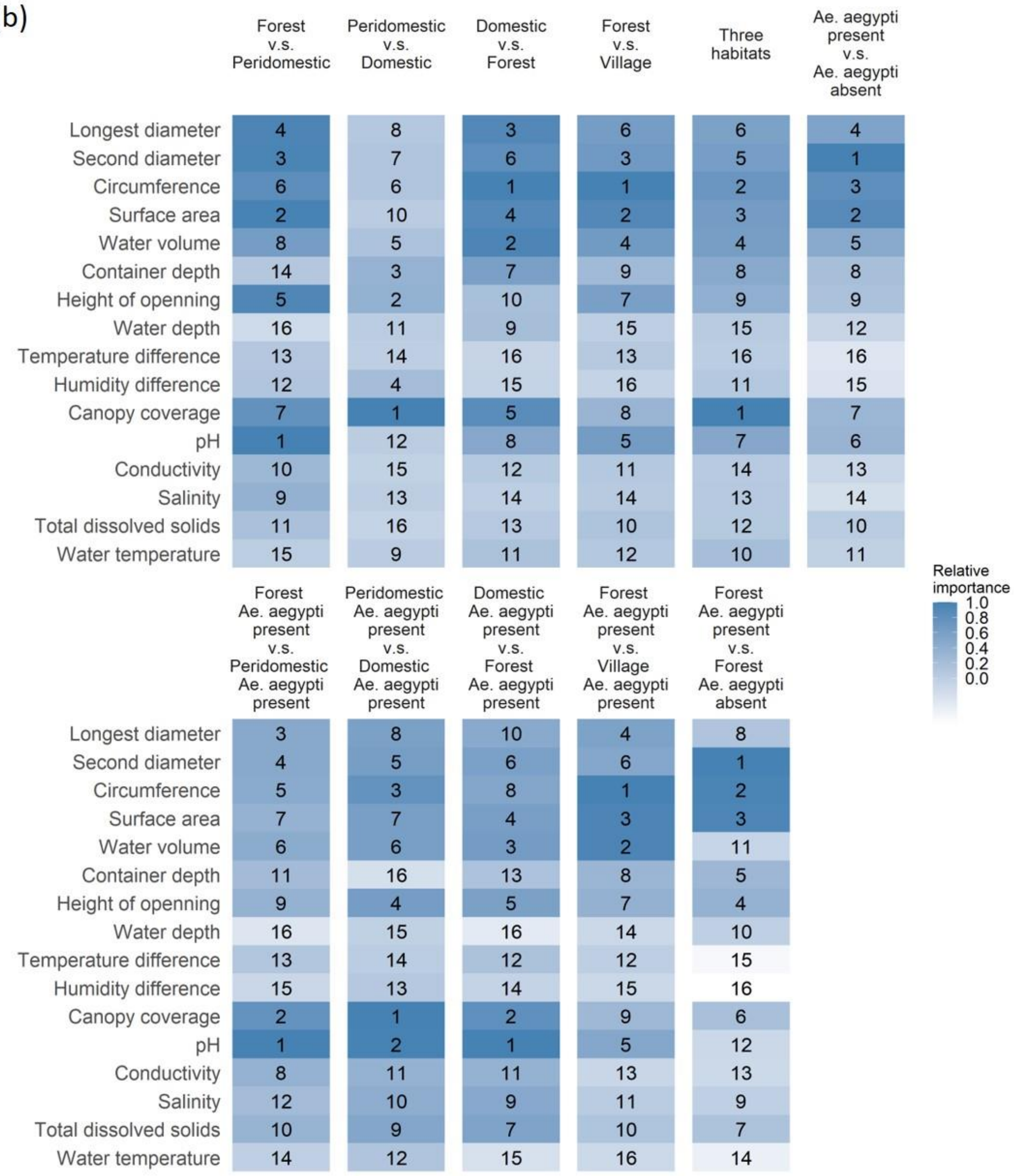

Forest v.s. Peridomestic
Peridomestic v.s. Domestic
Domestic v.s. Forest
Forest v.s. Village
Three habitats
Ae. aegypti present v.s. Ae. aegypti absent
Longest diameter 4 8 3 6 6 4
Second diameter 3 7 6 3 5 1
Circumference 6 6 1 1 2 3
Surface area 2 10 4 2 3 2
Water volume 8 5 2 4 4 5
Container depth 14 3 7 9 8 8
Height of openning 5 2 10 7 9 9
Water depth 16 11 9 15 15 12
Temperature difference 13 14 16 13 16 16
Humidity difference 12 4 15 16 11 15
Canopy coverage 7 1 5 8 1 7
pH 1 12 8 5 7 6
Conductivity 10 15 12 11 14 13
Salinity 9 13 14 14 13 14
Total dissolved solids 11 16 13 10 12 10
Water temperature 15 9 11 12 10 11
Relative importance
1.0
0.8
0.6
0.4
0.2
0.0
Forest Ae. aegypti present v.s. Peridomestic Ae. aegypti present
Peridomestic Ae. aegypti present v.s. Domestic Ae. aegypti present
Domestic Ae. aegypti present v.s. Forest Ae. aegypti present
Forest Ae. aegypti present v.s. Village Ae. aegypti present
Forest Ae. aegypti present v.s. Forest Ae. aegypti absent
Longest diameter 3 8 10 4 8
Second diameter 4 5 6 6 1
Circumference 5 3 8 1 2
Surface area 7 7 4 3 3
Water volume 6 6 3 2 11
Container depth 11 16 13 8 5
Height of openning 9 4 5 7 4
Water depth 16 15 16 14 10
Temperature difference 13 14 12 12 15
Humidity difference 15 13 14 15 16
Canopy coverage 2 1 2 9 6
pH 1 2 1 5 12
Conductivity 8 11 11 13 13
Salinity 12 10 9 11 9
Total dissolved solids 10 9 7 10 7
Water temperature 14 12 15 16 14

**Figure S3.** The importance of each physical variables in differentiating different comparisons between habitats or oviposition site groups in (a) La Lopé and (b) Rabai. Each row represents one variable, and each column describes the comparison, which is labeled above the figures. The color of each cell quantifies the relative importance, which is estimated by random forest in R. The absolute importance measures of all variables in each comparison were scaled to proportions of the variable that shows the highest importance measure. The numbers in each cell indicate the rank of the variables.

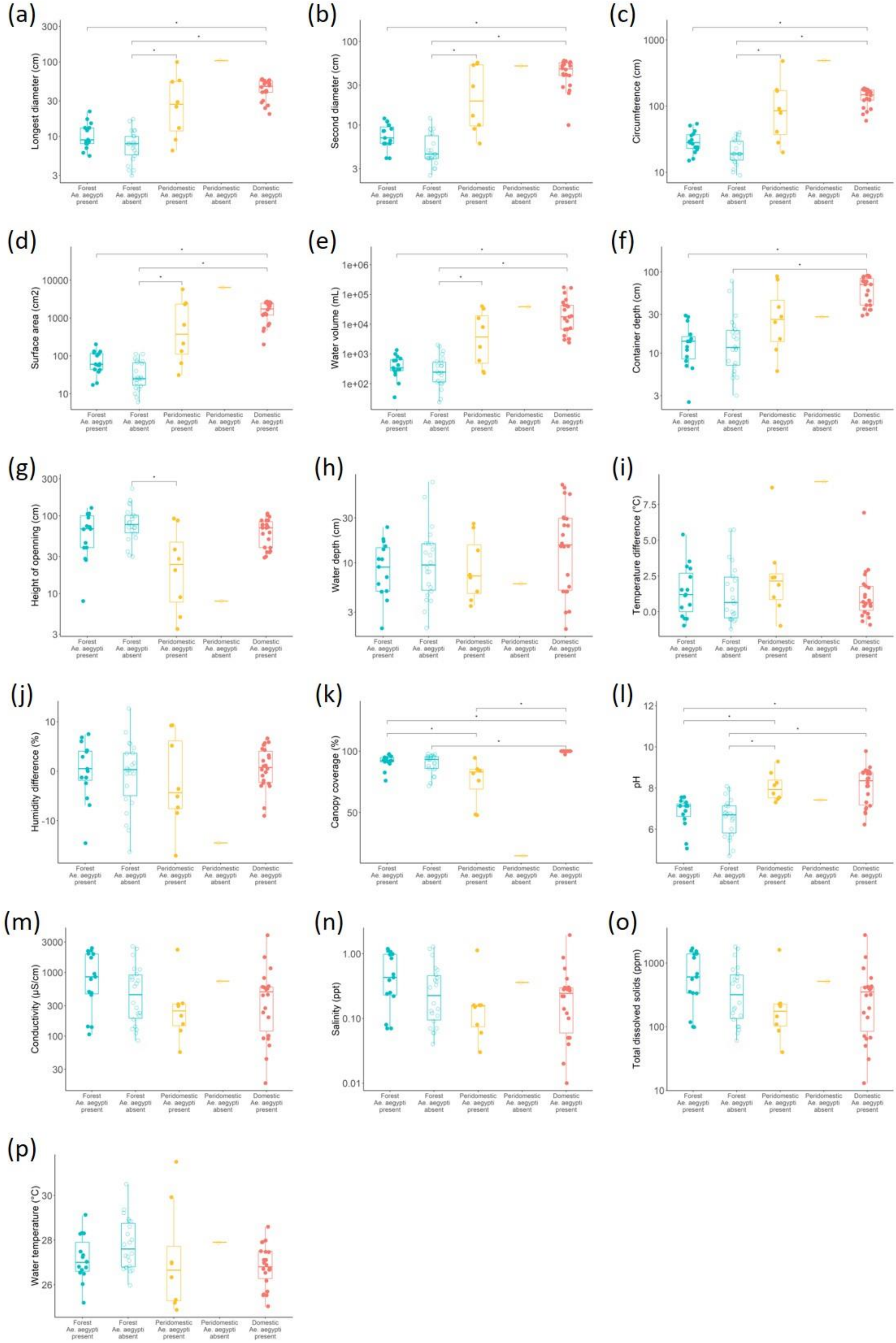
(a)
Longest diameter (cm)
(b)
Second diameter (cm)
(c)
Circumference (cm)
(d)
Surface area (cm2)
(e)
Water volume (mL)
(f)
Container depth (cm)
(g)
Height of openning (cm)
(h)
Water depth (cm)
(i)
Temperature difference (°C)
(j)
Humidity difference (%)
(k)
Canopy coverage (%)
(l)
pH
(m)
Conductivity (µS/cm)
(n)
Salinity (ppt)
(o)
Total dissolved solids (ppm)
(p)
Water temperature (°C)
Forest Ae. aegypti present
Forest Ae. aegypti absent
Peridomestic Ae. aegypti present
Peridomestic Ae. aegypti absent
Domestic Ae. aegypti present

**Figure S4.** Comparison of individual physical variables between oviposition site groups in Rabai. Each point represents a single oviposition site, and the boxplots show the minimum, 25% quartile, median, 75% quartile, and maximum of all values. The colors and shapes are as in Figure 2. Differences between groups were tested using pairwise Wilcoxon rank sum test with *Holm* multiple comparison correction (*: $p < 0.05$, Table S5 and S6). Only significant comparisons are labeled.

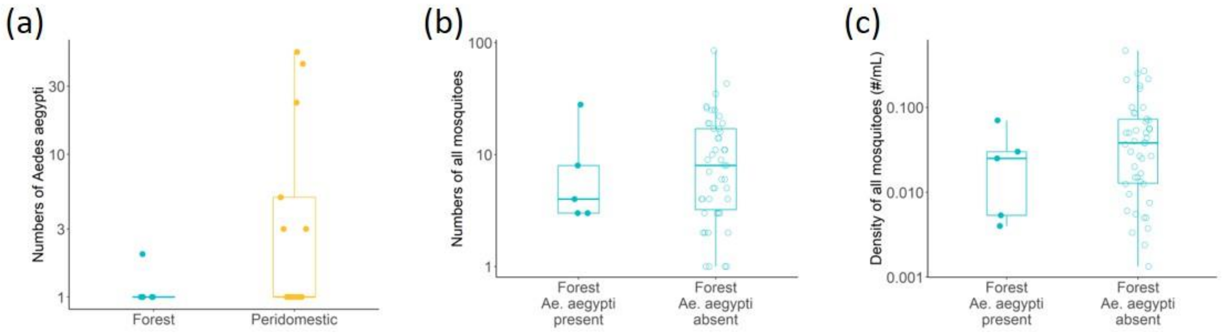


**Figure S5.** Comparison of (a) number of *Ae. aegypti*, (b) number of mosquitoes of all species, and (c) density of mosquitoes of all species in La Lopé. Mosquito species other than *Ae. aegypti* were only screened in the forest habitat, so in (c), we only compared forest oviposition sites present vs. absent of Ae. aegypti. Each point represents a single oviposition site, and the boxplots show the minimum, 25% quartile, median, 75% quartile, and maximum of all values. The colors and shapes are as in Figure 2. Differences of mosquito numbers between groups (a and b) were examined by negative-binomial models tests (*: $p < 0.05$), while differences of mosquito density (c) between groups were tested using pairwise Wilcoxon rank sum. Only significant comparisons are labeled.

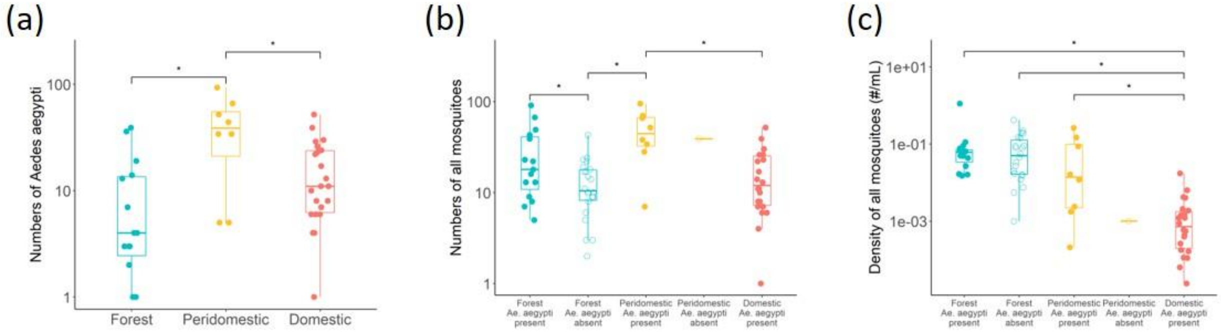


**Figure S6.** Comparison of (a) number of *Ae. aegypti*, (b) number of mosquitoes of all species, and (c) density of mosquitoes of all species in Rabai oviposition site between habitats or oviposition site groups. Each point represents a single oviposition site, and the boxplots show the minimum, 25% quartile, median, 75% quartile, and maximum of all values. The colors and shapes are as in Figure 2. Differences of mosquito numbers between groups (a and b) were examined by negative-binomial models, while differences of mosquito density (c) between groups were tested using pairwise Wilcoxon rank sum test with *Holm* multiple comparison corrections (*: $p < 0.05$, Table S5 and S6). Only significant comparisons are labeled.

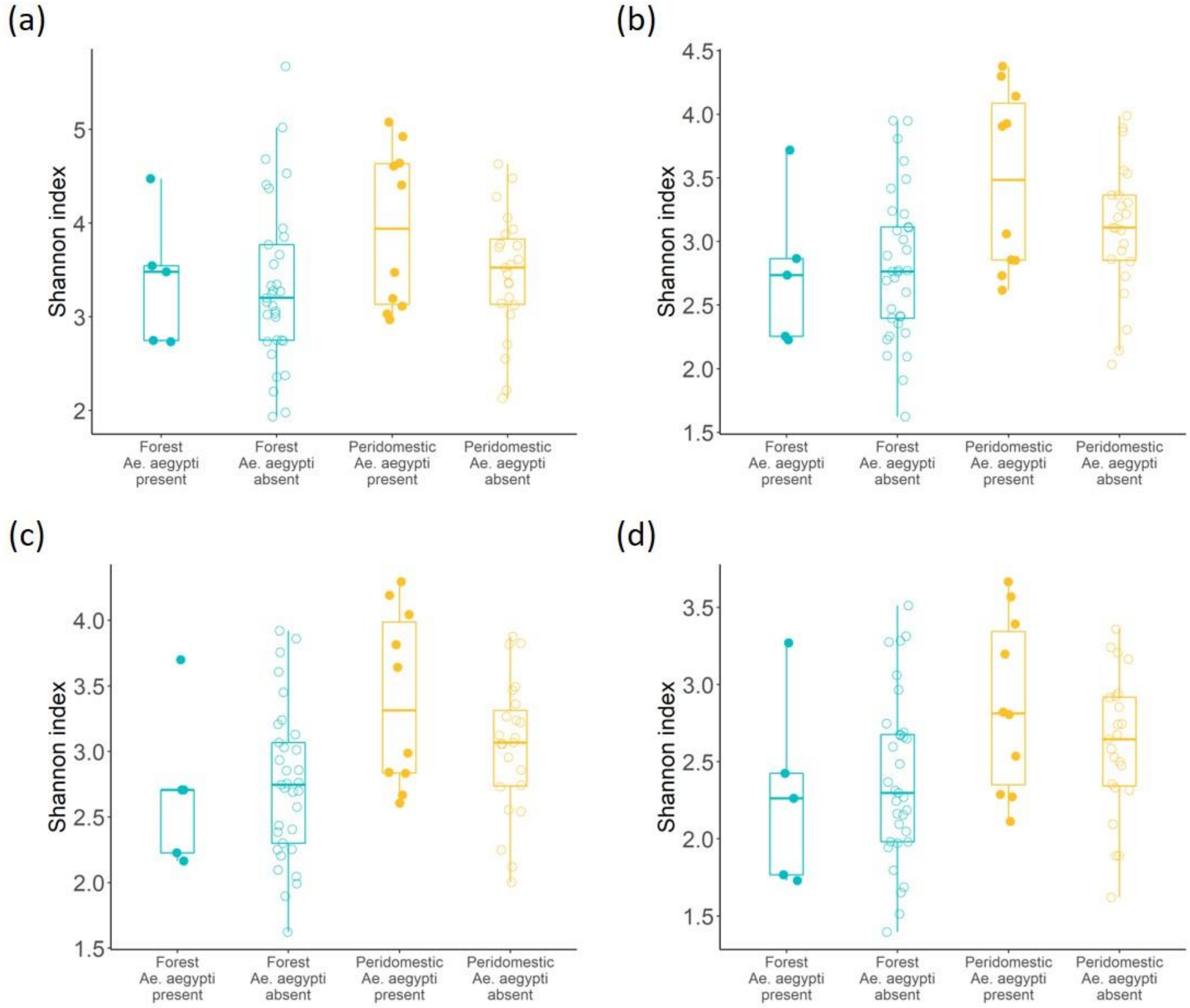


**Figure S7.** Comparison of the Shannon index of the bacterial community in La Lopé oviposition sites at different taxonomic levels: (a) ASVs, (b) Species, (c) Genus, and (d) Family. Each point represents a single oviposition site, and the boxplots show the minimum, 25% quartile, median, 75% quartile, and maximum of all values. The colors and shapes are as in Figure 2. Differences between groups were tested using pairwise Wilcoxon rank sum test with *Holm* multiple comparison corrections. No significant difference was found in any tests (Table S3 and S4).

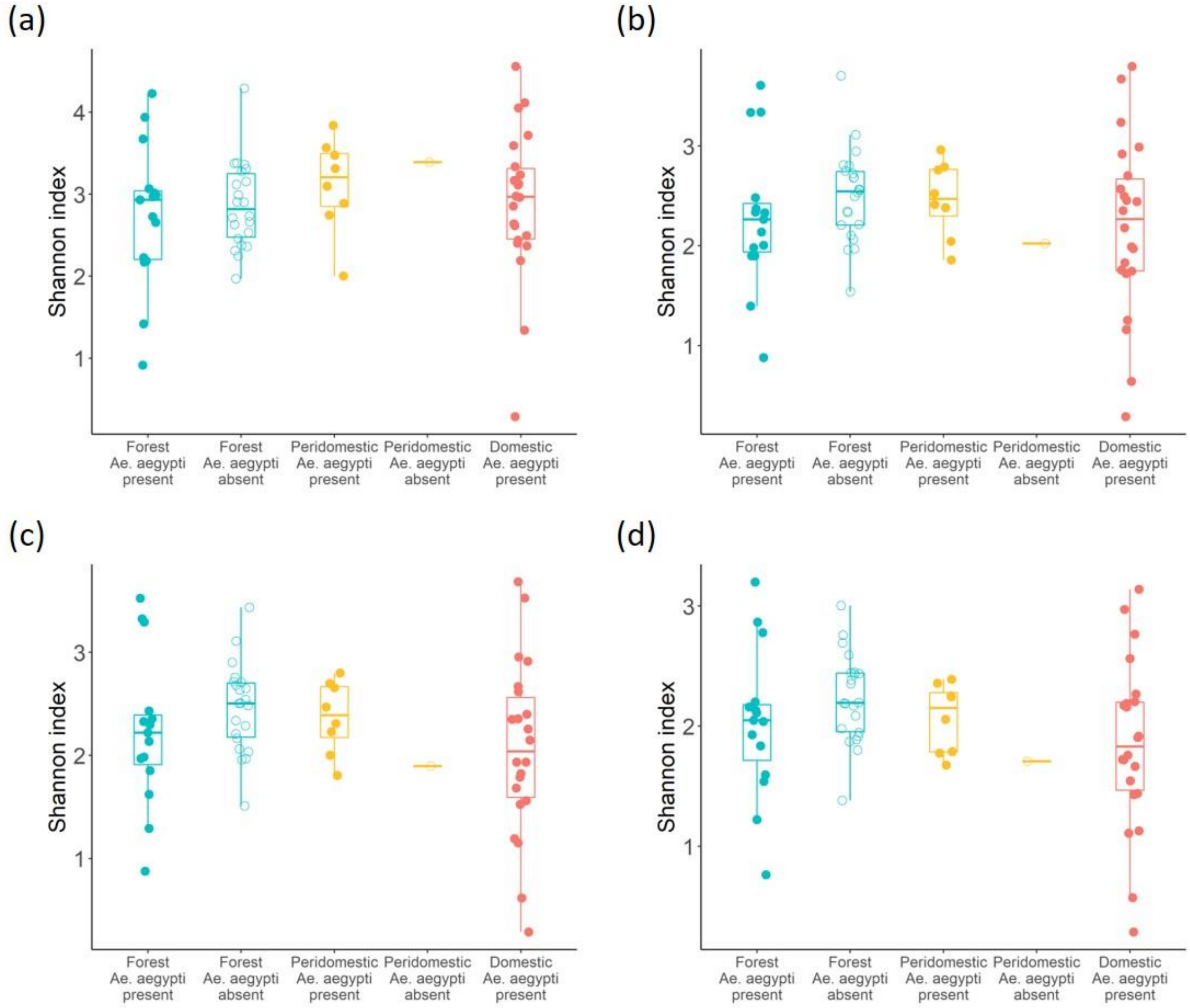


**Figure S8.** Comparison of the Shannon index of the bacterial community in Rabai oviposition sites at different taxonomic levels: (a) ASVs, (b) Species, (c) Genus, and (d) Family. Each point represents a single oviposition site, and the boxplots show the minimum, 25% quartile, median, 75% quartile, and maximum of all values. The colors and shapes are as in Figure 2. Differences between groups were tested using pairwise Wilcoxon rank sum test with *Holm* multiple comparison corrections. No significant difference was found in any tests (Table S5 and S6).

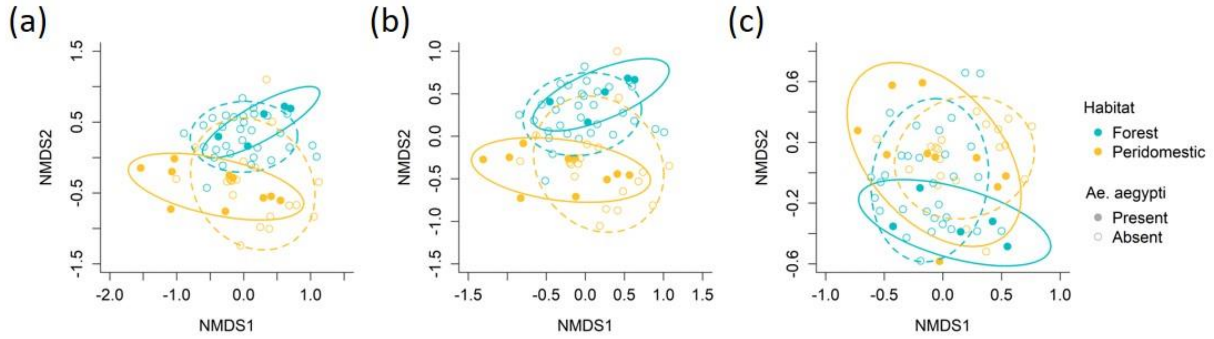


**Figure S9.** NMDS analysis of bacterial community compositions in La Lopé oviposition site at (a) Species, (b) Genus, and (c) Family level. Each point represents an oviposition site. The color and shape of points, as well as the ellipses, are the same as in Figure 2.

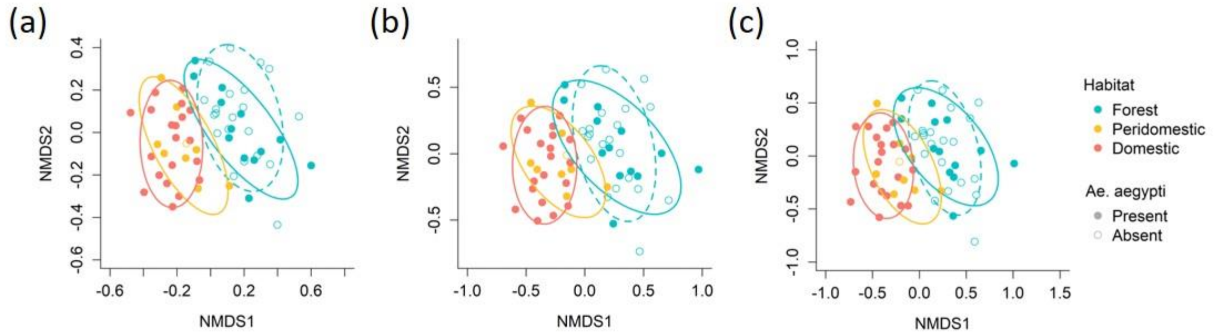


**Figure S10.** NMDS analysis of bacterial community compositions in Rabai oviposition site at (a) Species, (b) Genus, and (c) Family level. Each point represents an oviposition site. The color and shape of points, as well as the ellipses, are the same as in Figure 2.

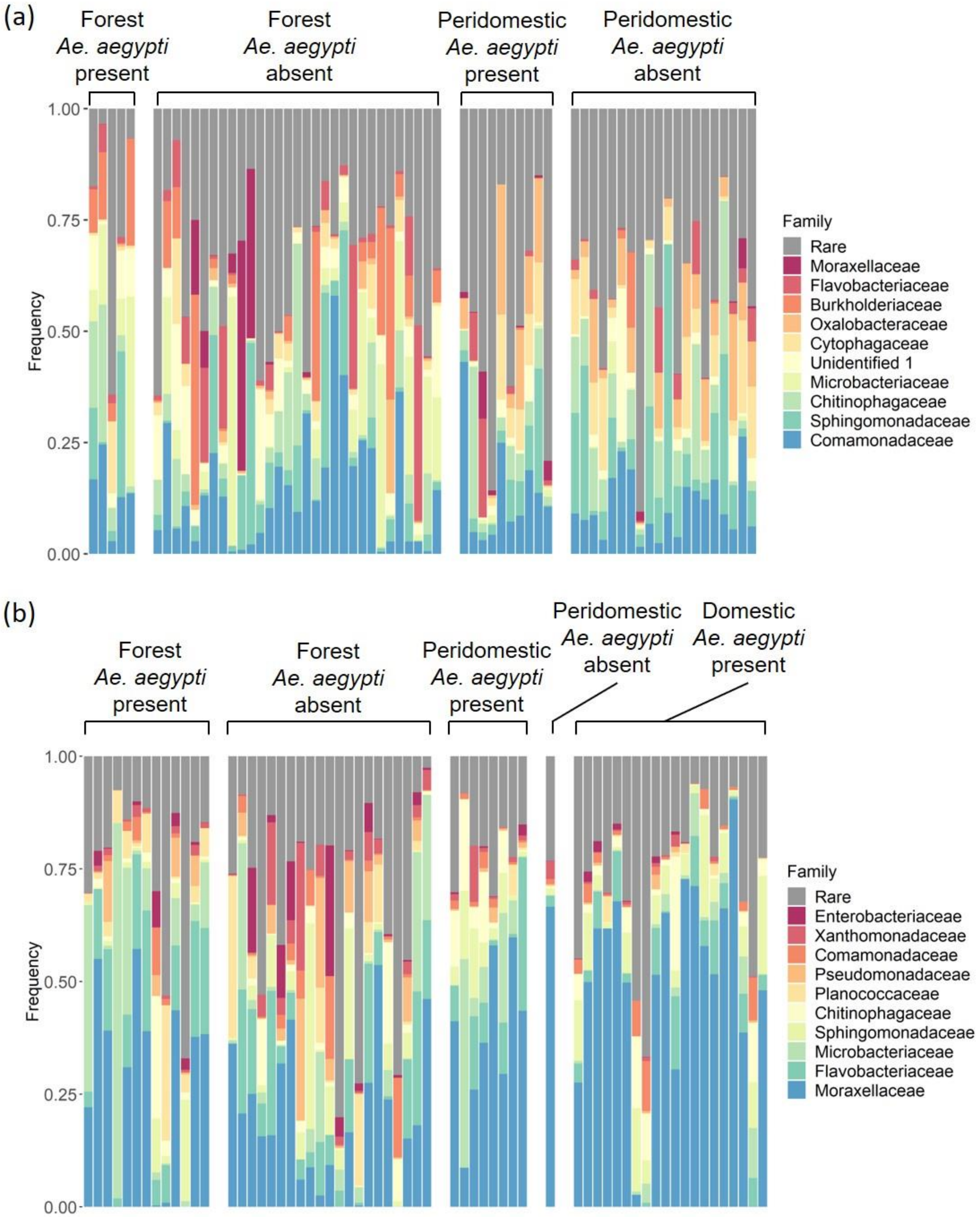


**Figure S11.** Frequency of the top ten bacterial Families in oviposition site in (a) La Lopé and (b) Rabai oviposition sites. Each bar represents an oviposition site, and the length of each color represents the proportion of the corresponding Family in the site. Other bacterial Families are grouped in the “Rare” category, which shows as gray in the bar plots. Samples are grouped by oviposition site groups, which are labeled above the bar plots.

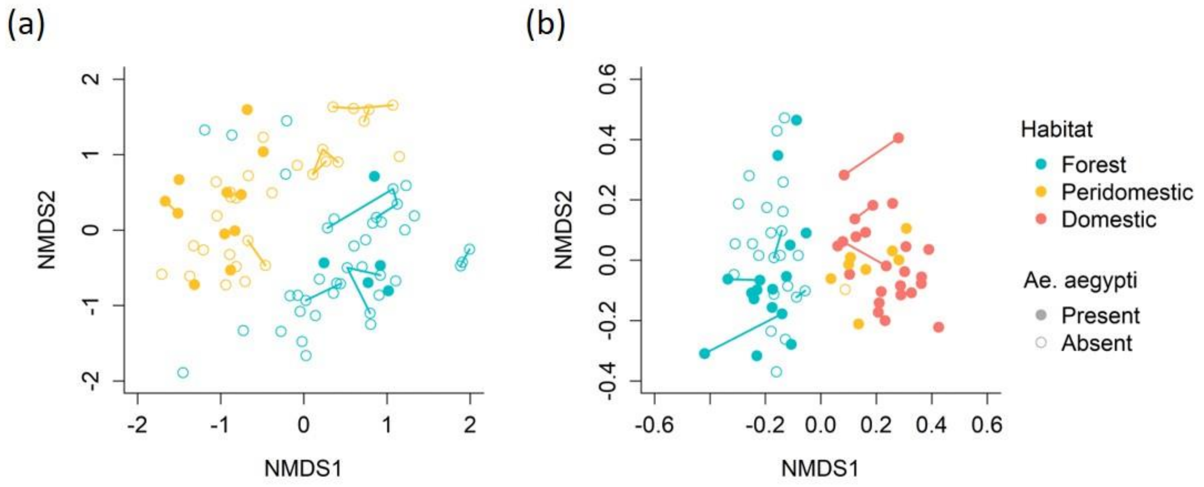


**Figure S12.** Temporal variation of bacterial community compositions at the ASV level in oviposition sites in (a) La Lopé and (b) Rabai. Each point represents an oviposition site. The color and shape of points, as well as the ellipses, are the same as in Figure 2. Bacterial samples collected from the same oviposition sites at different times are linked with segments.

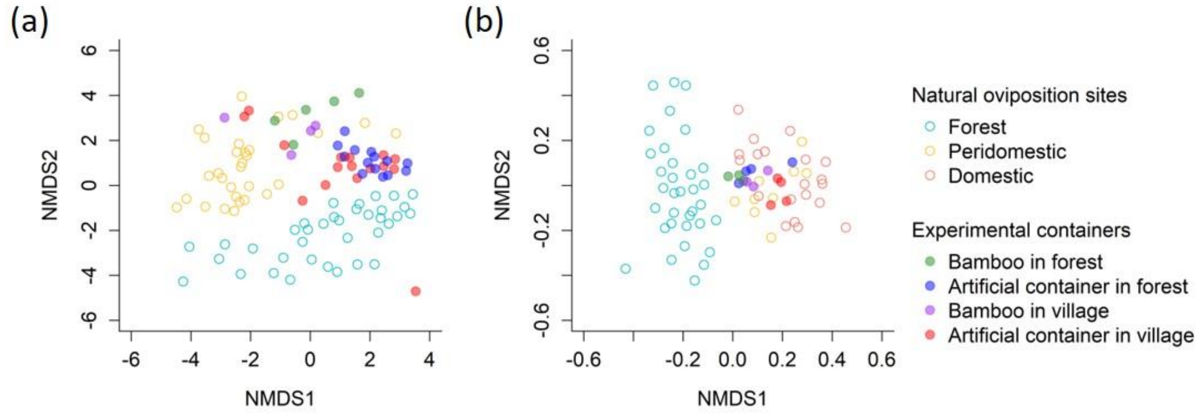


**Figure S13.** Bacterial community compositions of the containers used in the field oviposition choice experiments in (a) La Lopé and (b) Rabai. The NMDS was performed at the ASV level. Each point represents an oviposition site. Hollow points represent natural oviposition sites, and solid points represent experimental containers. The two types of experimental containers in the two habitats were differentiated by different colors.

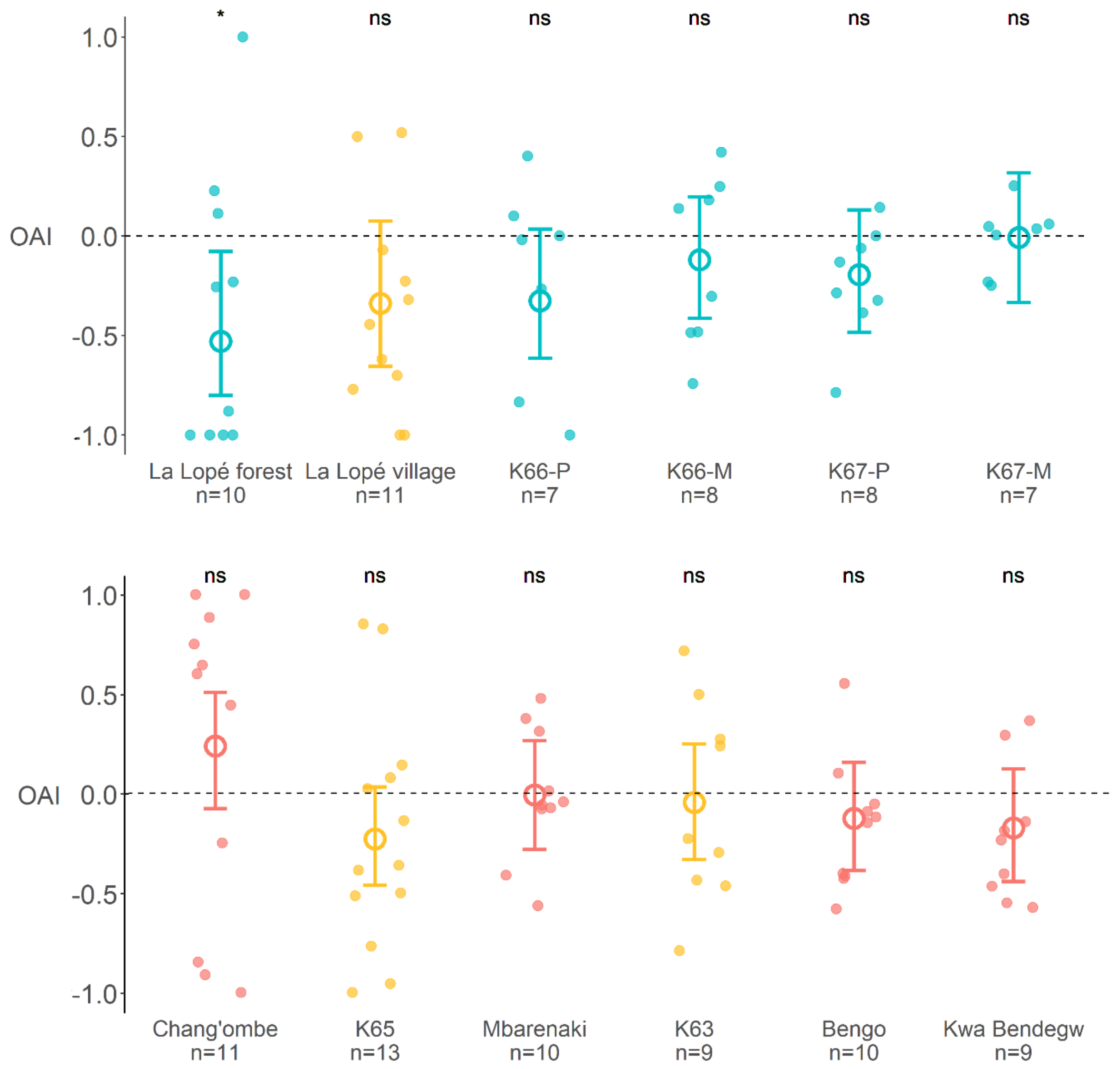


**Figure S14.** Results of laboratory oviposition assay on forest vs. village bacteria culture summarized by colonies. The details of the two bacterial cultures were described in Table S2. Each point represents one cage with five gravid females, and the color indicates the habitat where the colony originated. Higher OAI indicates preference for forest bacteria cultures. K66-P and K66-M represents the two copies of the same forest colony maintained at Yale and Princeton University. A beta-binomial model was used to test differential preference among colonies, which results in no significant colony effects. The model also estimated the mean OAIs and the 95% confidence intervals of all colonies, indicated by the hollow cycles and the error bars. The asterisks and ‘ns’ above each colony indicates whether the 95% CI excludes zero.

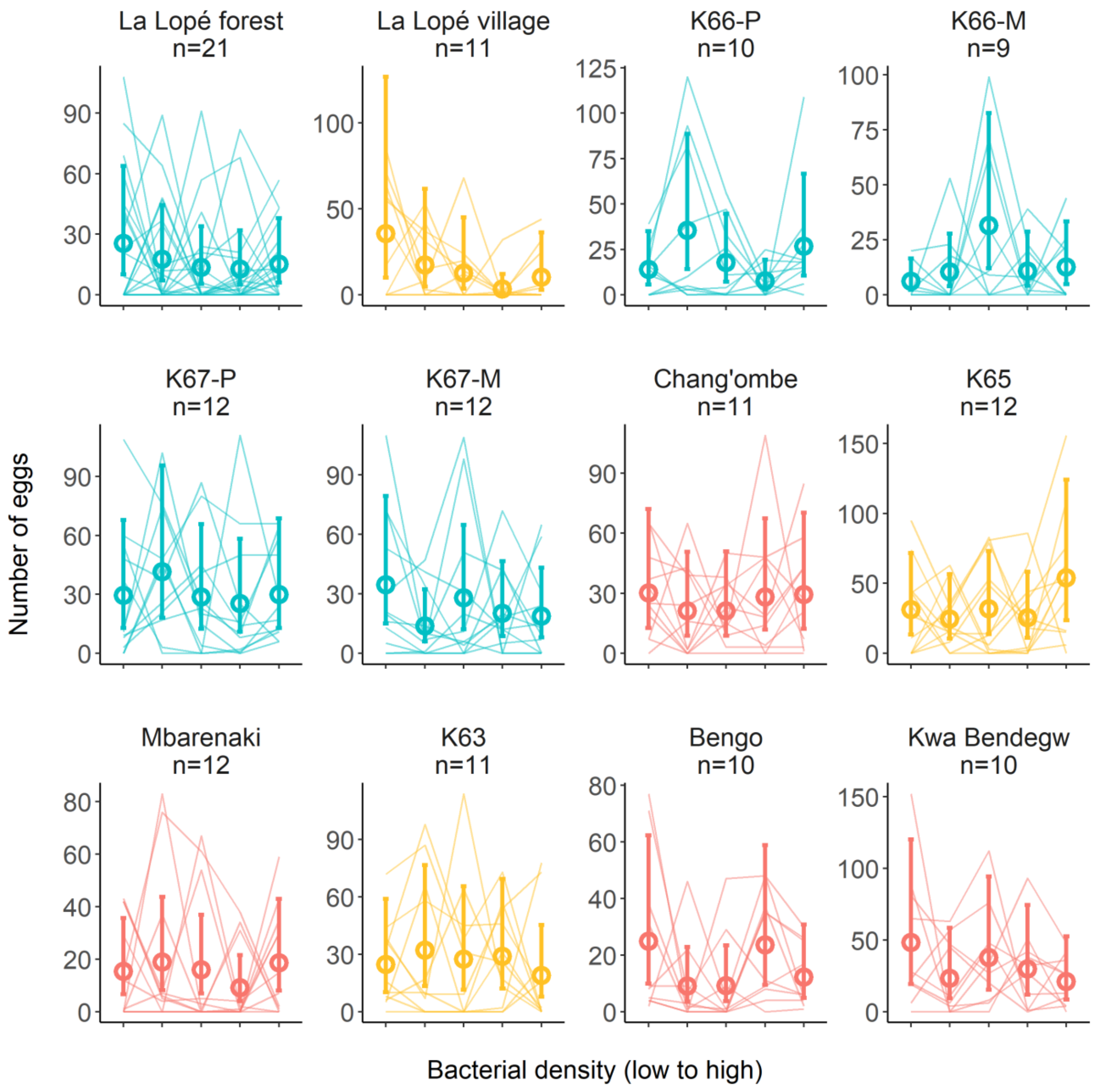


**Figure S15.** Results of laboratory oviposition assay on bacterial density summarized by colonies. Five cups were provided in each cage with increasing bacterial density at 0, $2 \times 10^5$, $1 \times 10^6$, $5 \times 10^6$, $2.5 \times 10^7$ cells/mL (details in Table S2), which correspond to the five columns in each subplot. Each line connects the five egg counts in one cage. Colors represent the habitats from where the colonies came. A negative-binomial model was used to fit the results of each oviposition assay. The model estimates the mean number of eggs in each bacterial density and a 95% confidence interval, which are shown by the hollow cycles and the error bars, respectively.
